# Learning is a fundamental source of individuality

**DOI:** 10.1101/2024.08.30.610528

**Authors:** Riddha Manna, Johanni Brea, Gonçalo Vasconcelos Braga, Alireza Modirshanechi, Ivan Tomić, Ana Marija Jakšić

## Abstract

Learning and memory are essential components of our individuality. While it is established that behaviour can vary across genetically identical individuals, it remains unknown how much of this variation stems from momentary experience during learning compared to genetics and its past interactions with the environment. To address this, we measured behaviour in thousands of flies from 90 genetic backgrounds while they performed tasks in conditions that either did or did not require learning. Flies that were genetically identical, raised under the same conditions and tested simultaneously in the same environment persistently modified the extent of expressed individuality when they could learn. This learning-induced residual expression of individuality and its dynamics were subdued or absent in innate, learning-independent behaviours. We could quantify and then recreate this phenomenon in computer simulations. The emergence of *in silico* behavioural individuality was most consistent with the individuality of real flies once we enabled reinforced learning in simulated agents. Moreover, we showed that minor differences in initial conditions of the experiment can exacerbate the expression of individuality within a genotype in a learning-dependent manner. Our results establish that besides the classical G x E interactions shared between individuals in the past, learning from individual momentary experience further extends the expression of individuality.

## Introduction

Behavioural individuality is an inevitable, fundamental aspect of life. Even identical twins who share a genome and life history eventually diverge into distinct individuals ***Koellinger and Harden (2018)***; ***Martin (2005)***; ***Linneweber et al. (2020)***. In every individual, behaviour is shaped by fixed, genetic factors ***Dudai et al. (1976)***; ***Partridge and Sgrò (1998)***; ***Ayroles et al. (2015)***; ***Harden (2023)*** and by variable environmental events throughout lifetime, which may be stochastic and can occur at the molecular, cellular, organismal and even population scales ***Anderson et al. (2017)***; ***Honegger and de Bivort (2018)***; ***Linneweber et al. (2020)***; ***Graham (2021)***; ***Laskowski et al. (2022)***; ***Harden (2023)***; Thomas F. ***Mathejczyk et al. (2023)***; ***Maloney et al. (2024)***. These classical sources of individuality have been studied extensively in the context of development ***Korner (1971)***; ***Boogert et al. (2018)***; ***Linneweber et al. (2020)***; ***Laskowski et al. (2022)***. Modifications to individuality after development are less well understood. Behaviour is known to change on average during adult life. The change occurs either through learning and memory, or through learning-independent processes, as an adaptation to environments experienced during life ***Buchanan et al. (2015)***; ***Xu et al. (2016)***; ***Boogert et al. (2018)***; ***Mollá-Albaladejo and Sánchez-Alcañiz (2021)***; ***Smith et al. (2022)***; ***Modi et al. (2023)***; ***Maloney et al. (2024)***. Intriguingly, the ability to learn and remember is itself known to vary across individuals ***Turner (1907***, ***1911***); ***Kandel and Hawkins (1992)***; ***Boogert et al. (2018)***; ***Smith et al. (2022)***, genotypes ***Nepoux et al. (2015)***; ***Williams-Simon et al. (2019)***, environments ***Tosh and Brogan (2017)***; ***Wang et al. (2018)***; ***Lin et al. (2020)***, or even maternal status ***Mao et al. (2018)***; ***Luo et al. (2017)***; ***Álvarez Quintero and Kim (2024)***.

Traditionally, the cause of variability in learning ability has been viewed the same way as variability in any other phenotype - as an average outcome of genotype x environment (G x E) interactions measured within a genotype. However, individual learning is a critical individual manifestation of G x E, where the interaction occurs not only in the past of the individual (prior to measuring behaviour), but also at the moment the environment is experienced through behaviour. Importantly, this momentary individual experience of the environment during learning can uniquely impact not only the behaviour but also what the future experience will be for only that particular individual. This particular contribution of individual learning on the expression of individuality has not been studied extensively because the measurement of G x E for learning-dependent behaviour cannot be easily averaged over other individuals or over time.

The challenge in studying the role of individual learning experience on individual behaviour lies precisely in its non-ergodic nature: momentary experience can cause learning, and learning can change future experience. Learning is a dynamic product of genetics and environment (Figure 1A). To study it, it is necessary to experimentally control genetics and environment over generations, and ensure that learning experience is temporally parallel across tested individuals. Most previous studies have for this reason focused on individuality in spontaneous behaviours that are insensitive to momentary experience and independent of learning. Such behaviours can be measured repeatedly and can be averaged over time for a single individual ***Mery and Burns (2010)***; ***Chen and Sokolowski (2022)***; ***Korner (1971)***; ***de Bivort et al. (2022)***; ***Linneweber et al. (2020)***; ***Kempermann et al. (2022)***; ***Mollá-Albaladejo and Sánchez-Alcañiz (2021)***; ***Sridhar et al. (2021)***; Thomas F. ***Mathejczyk et al. (2023)***; ***Maloney et al. (2024)***. In the few studies that do address learning behaviour, the influence of both genetics and individual environment is never accounted for simultaneously, mainly due to technical limitations of executing such an experimental design ***Arican et al. (2020)***; ***Finke et al. (2021)***; ***Smith et al. (2018)***; ***Galsworthy et al. (2005)***; ***Williams-Simon et al. (2019)***; ***Chittka et al. (2003)***; ***Nouvian and Galizia (2019)***; ***Giurfa et al. (2001)***; ***Smith et al. (2022)***. Namely, the lack of simultaneous measurements of behaviour prevents distinguishing the effects of momentary from past experience on behaviour.

**Figure 1.**
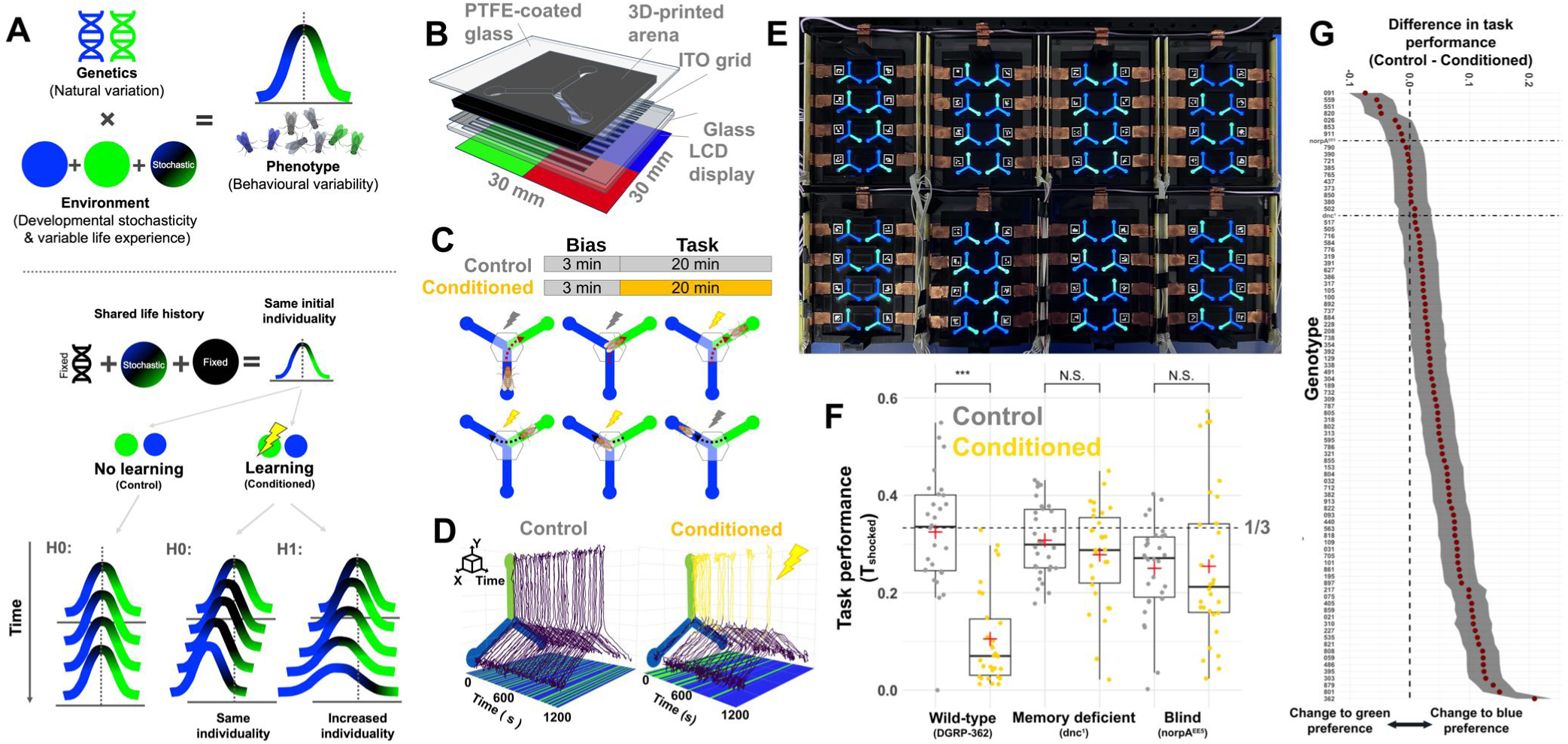
Capturing sources of individual learning-dependent behaviour. (A) Variation in behaviour between individuals can be caused by variation in genetic background, variation in environment (which can be fixed and shared between individuals or stochastically individual), and genotype x environment interactions. We can experimentally fix the genotype x environment interaction in the past and thus control the shared extent of individuality at the beginning of the task. When task behaviour is measured in parallel across individuals, we expect that the extent of behavioural variability within genotype will correlate between spontaneous and learning-dependent behaviour, while the genotype means may change (H0). Alternatively, learning may change the extent of individuality as well as genotype means over time (H1). (B) Schematic of the Y-maze design. (C) During the course of the assay each fly is first tested for its individual behavioural biases. Then, the task part of the assay is initiated for all flies simultaneously. Control flies are not shocked when choosing green colour during the task, while conditioned flies are. Shock delivery and shock removal timing in conditioned task for a bad decision (blue to green) and after a bad decision (green to blue) is depicted below. (When switching from blue to blue, no shock is ever applied.) (D) Representative tracks of two flies traversing a Y-maze in control and conditioned task. Lines on the bottom plane represent colour occupancy over time. (E) Top-down view of the 64-maze multiplexed platform. (F) Task performance of a wild-type genotype compared to the memory- and learning-deficient mutant (*dnc1*) and the visually impaired mutant (*norpAEE5*). Points are individual fly measurements. Mean and medians are represented as bar and red plus. Dashed line represents chance level T_shocked_ = 1/3. (G) Difference in mean task performance between conditioned and control flies in 90 genotypes. Grey shading is SE. Vertical dashed line indicates no change in average performance upon conditioning. Horizontal dashed lines indicate mean values of memory- and vision-deficient genotypes (*dnc1* and *norpAEE5*, horizontal). **Figure 1—figure supplement 1. Operant conditioning paradigms in a Y-maze.** **Figure 1—figure supplement 2. Multiday-multiparadigm operant conditioning in wild-type (DGRP-362) genotype and memory deficient genotype (dnc^1^).**

In genetically identical individuals who share both the same past as well as momentary experiences, it is expected that the variance in behaviour across individuals will correlate between spontaneous behaviour and learned behaviour (Figure 1A). This is because, when the environment is experienced by all individuals in parallel throughout the entire life, including during learning, individual learning experiences will effectively constitute the same genotype x environment interaction, one that is fully shared among individuals and can now be averaged across them. Of course, stochasticity of the environment over time is still expected to affect individuals and produce variation, but with temporal coincidence, it will be uniformly affecting flies that do and do not learn. The average behaviour may change between learning and non-learning flies, as well as the variance, but the shape of the distribution is expected to stay the same. Alternatively, stochasticity of the learning process across individuals could diversify behaviour over time. However, even in this case, the change in expressed individuality should be similar across genotypes.

In this study, we develop an experimental setup to test this expectation. We leverage the Drosophila model system, where we could achieve extreme uniformity in experienced environment during learning in thousands of isogenic individuals (Table 1) across 90 different genetic backgrounds. We analysed, modelled, and then computationally recreated the time-resolved experiences of individual flies as they made nearly half a million decisions, while they were learning or behaving spontaneously. This allowed us to partition and quantify the independent contributions of all measurable sources of individuality on variation in behaviour, including genetics, shared environment during development and life, and momentary environmental experience during behaviour (Figure 1A). Our findings showed that behavioural diversity across individuals changes due to individual learning, even when genetic, past and momentary environmental factors are held constant, highlighting the stochastic processes during learning as fundamental to individuality as developmental stochasticity.

**Table 1.** Distribution of individuals - place learning green in wild-type, memory mutant and blind mutant genotype experiment.

| Genotype | Replicate 1 (32) |  |  |  | Replicate 2 (32) |  |  |  | Total (64) |
| --- | --- | --- | --- | --- | --- | --- | --- | --- | --- |
|  | Male (16) |  | Female (16) |  | Male (16) |  | Female (16) |  |  |
|  | Control (8) | Conditioned (8) | Control (8) | Conditioned (8) | Control (8) | Conditioned (8) | Control (8) | Conditioned (8) |  |
| Label: | noShock_male_rep1 (8) | shock_male_rep1 (8) | noShock_female_rep1 (8) | shock_female_rep1 (8) | noShock_male_rep2 (8) | shock_male_rep2 (8) | noShock_female_rep2 (8) | shock_female_rep2 (8) |  |
| DGRP362 | 8 | 8 | 6 | 7 | 7 | 7 | 7 | 8 | 58 |
| dnc<sup>1</sup> | 8 | 8 | 8 | 7 | 8 | 8 | 6 | 7 | 60 |
| NorpA<sup>ee5</sup> | 8 | 8 | 8 | 6 | 8 | 8 | 6 | 8 | 60 |
| Place learning green |  |  |  |  |  |  |  |  |  |
| Number of flies used to analyse the data after all the filtering steps to filter out flies for activity, injury and their distribution across replicates, sexes and conditioning. Number of flies placed in the arenas at the start of the assay are mentioned within brackets on top of each column. In this experiment, flies were shocked when entering the arm with green colour. That arm is always green, and the other two are blue. Thus, the fly has to learn to avoid this arm (this place) and/or the green colour. |  |  |  |  |  |  |  |  |  |

**Table 2.**
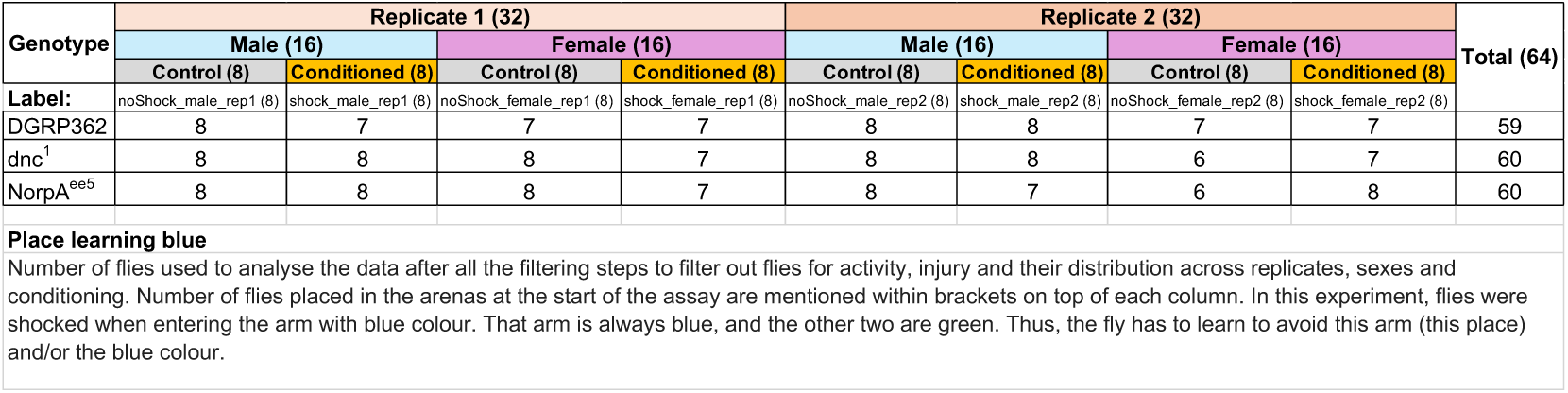
Distribution of individuals - place learning blue in subset of DGRP genotypes.

**Table 3.**
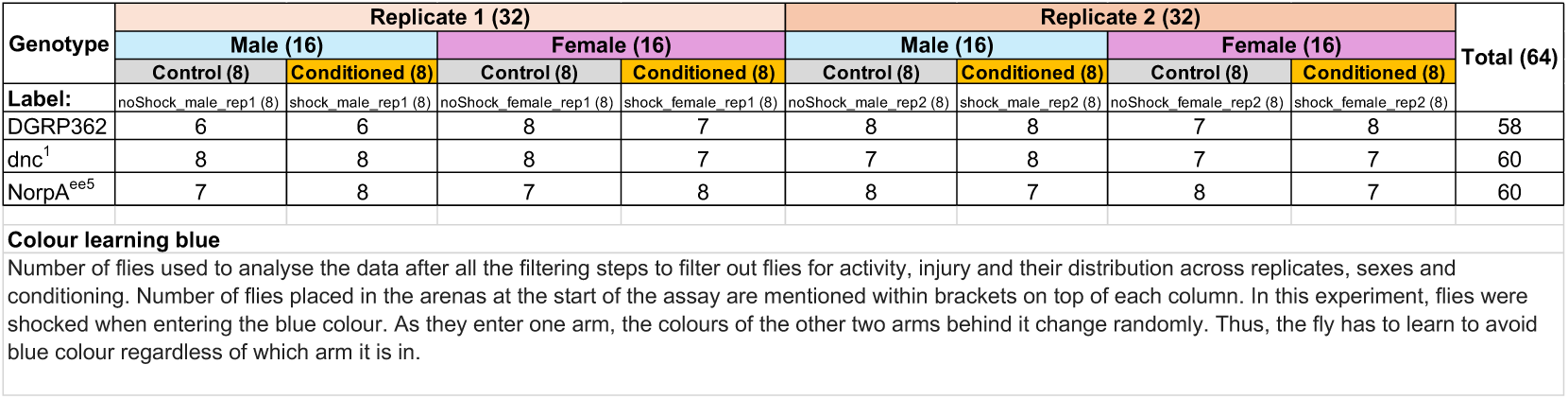
Distribution of individuals - colour learning green in wild-type, memory mutant and blind mutant genotype experiment.

**Table 4.**
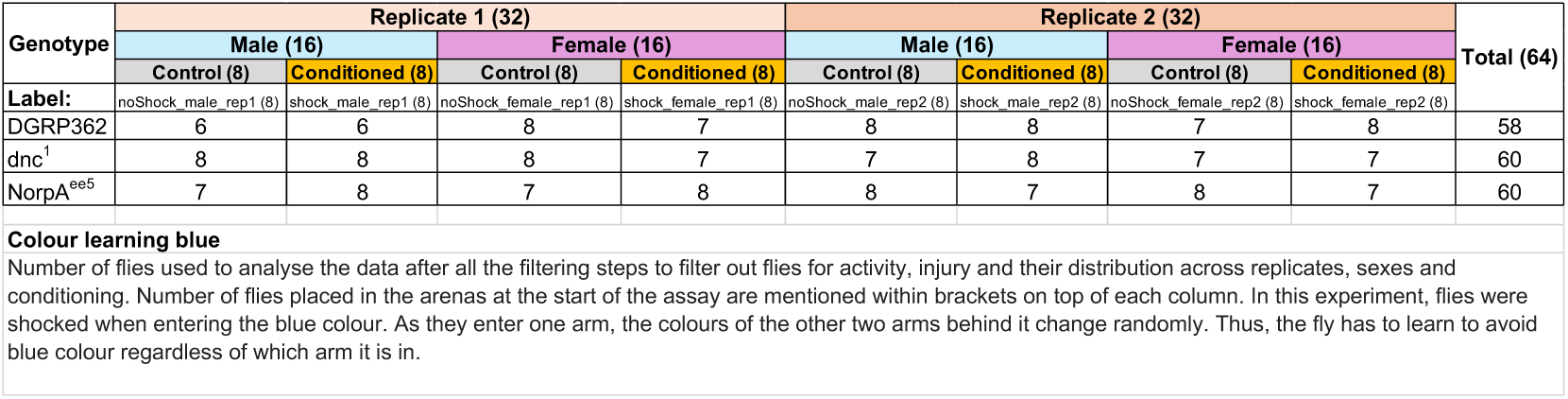
Distribution of individuals - colour learning blue in wild-type, memory mutant and blind mutant genotype experiment.

## Results

### Distributions of individual behaviours are altered by genotype and learning

Both genotype and environment affect behaviour. However, even in an experimentally controlled environment, a genotype also interacts with the inevitable environmental stochasticity - temporal fluctuations in temperature and atmospheric pressure, unexpected vibrations and air movement, or constantly changing molecular environments in the cell. Such small stochastic changes in the environment can add up. They primarily affect individuals during development, resulting in the ubiquitously observed normally distributed variance of measured phenotypes, even among genetically identical individuals raised in a shared environment ***Little (1933)***; ***Ayroles et al. (2015)***; ***Klingenberg (2019)***. This gaussian variance has been reliably used as a read-out of the extent of individuality expressed within a genotype ***Ayroles et al. (2015)***; ***de Bivort et al. (2022)***. We asked is the distribution of expressed behaviour within a genotype constant or altered in the presence of learning if all individuals shared the same life history and the same experience of the environment while the behaviour is tested.

We tested learning-dependent and spontaneous behaviours in all individuals of a genotype at the same time. This way every individual experienced the same environment (and its temporal stochasticity) during development and while behaving. To do so, we designed a platform that implements operant conditioning assay that can be parallelized across 64 freely behaving individual *Drosophila melanogaster* (Figure 1B, Figure 1E). The system takes input from real-time video tracking of individuals. The tracking is then used for closed-loop coupling of visual stimuli with a mild foot shock that is delivered to individual flies upon an undesired behaviour (Figure 1C, Figure 1D; Figure 1 - figure supplement 1A). The aversion to shock prompts the individual to learn to avoid it. In the assay, each individual fly freely moves in a Y-shaped arena (“Y-maze”). The floor of the arena is illuminated such that each arm of the maze projects a colour. The conditioned task for a fly is to reduce the proportion of time spent on a particular colour (or arm) that is associated with shock (*task performance* or T_shocked_, where smaller value represents better task performance).

We used this assay to first test the performance of known learning- and sensory-deficient flies ***Dudai et al. (1976)***; ***Harris and Stark (1977)*** and one wild-type genotype. We validated the assay using several learning paradigms, such as “green place learning”, “blue place learning”, “green colour learning” and “blue colour learning” (measured N=192, n=64 individuals per genotype and per paradigm, Table 1-4, Figure 1C-D,F, Figure 1- figure supplement 1A-B,D-E), and in a multiday experiment where paradigms were swapped each day for individual flies that were tracked over multiple days (Figure 1- figure supplement 2, Table 5). In the main experiment, we focused on the paradigm where flies are conditioned to avoid green colour that is always associated with the same place (”place learning green”) and tested 88 genetically diverse wild-type genotypes (N= 5632, n=64 individuals per genotype, Table 6) ***Mackay et al. (2012)*** (Figure 1F-G). We ensured strict multi-generational control for each genotype and their two biological replicates. Replicates were raised independently in separate vials, but otherwise at the same time and under same environmental conditions (Figure 2B, Table 6-8, Figure 2-figure supplement 1B-I). Individuals from each replicate (vial) were distributed evenly into conditioned and control group and their behaviour was tested simultaneously (Table 6, Table 8). Unlike in the conditioned group of flies, in the control group, the shock was never applied, allowing the flies to behave spontaneously, without a need to learn to avoid any colour (Figure 1C). Note that this also means that in the control group, task performance reflects innate colour preference, while in conditioned group it can reflect innate preference, learned colour avoidance, as well as shock avoidance. We tracked their momentary task-relevant behaviours, including their locomotor activity, handedness and ∼half a million individual decisions (i.e., switches from one arm in the Y-maze to another arm; N = 497450 decisions; Figure 1D).

**Figure 2.**
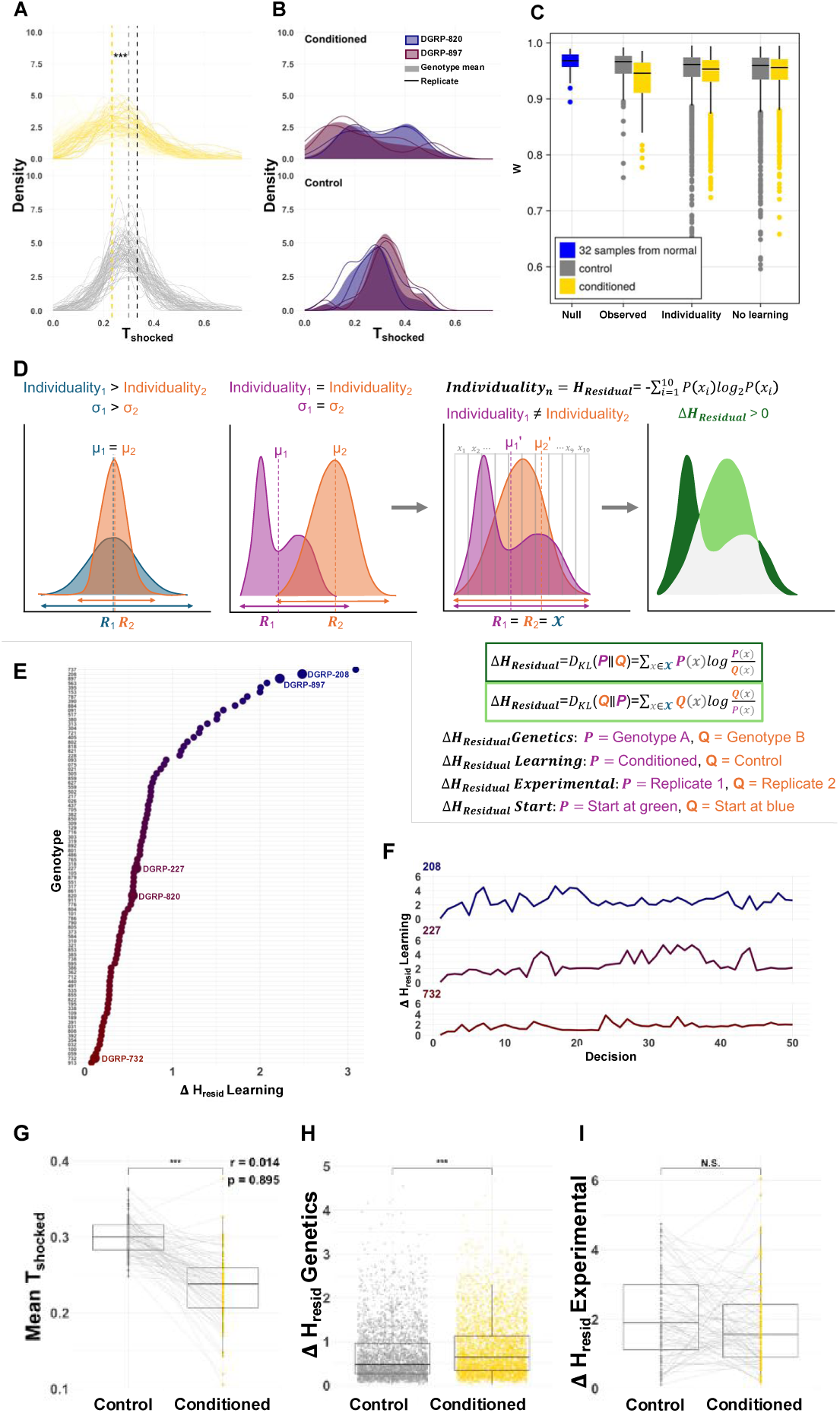
Residual individuality in task performance. (A) Distributions of individual task performance for the 88 wild-type genotypes, for conditioned (yellow) and control (gray) groups. Vertical lines show mean task performance. Chance expectation of T_shocked_ shown at 1/3 (black). (B) An example of individual task performance distributions in two genotypes (shaded red and blue). Replicates are shown as lines. (C) Shapiro-Wilk test for normality for distributions of observed individual task performance in 88 genotypes, and individual task performance of simulated flies that could or could not learn. In blue is a random sample from a normal distribution matching the observed task performance. (D) Measuring residual individuality. Difference in variance is commonly used to measure expressed individuality in behaviour when genetically identical individuals from two genotypes (e.g. blue and orange) are raised in identical environments. However, the shape of the distribution can change, even when variance remains the same (e.g. magenta and orange), suggesting still a change in the expression of individuality. By scaling the distribution to a fixed range (*R*=*X*) and calculating its entropy, residual individuality (H_resid_) represents the shape of the distribution, while change in residual indivduality (ΔH_resid_) captures the difference in shape (green) between two distributions, irrespective of the differences in their means or variances. E.g. when comparing blue distribution to orange, the change in residual individuality will be 0 because the shapes are the same. ΔH_resid_ is a measure of how difficult it is to approximate the shape of one distribution (*Q*) with the shape of the other (*P*). (E) Change in expressed residual individuality due to learning across genotypes. A few genotypes are highlighted to facilitate comparisons with other figures. (F) Divergence in residual individuality due to learning evolves with every decision and increases over time. Three representative genotypes are shown. (G) Mean task performance across genotypes (points). Lines connecting a genotype’s value in the two treatments indicate a significant correlation in mean task performance between conditioned and control groups. (H) Change in expressed residual individuality due to genetics indicates that the residual individuality is significantly more diverse between genotypes in conditioned flies. A point is a pairwise divergence between two genotypes. (I) Change in expressed residual individuality due to experimental procedures. There is no greater difference in residual <u>individuality between replicates in conditioned flies, compared to control. Point is a pairwise divergence between two replicates of a genotype.</u> **Figure 2—figure supplement 1. Density distributions of task performance.** **Figure 2—figure supplement 2. Task-relevant phenotypes.** **Figure 2—figure supplement 3. Correlations between individual behaviours.** **Figure 2—figure supplement 4. Correlations between genotype–averaged behaviours and shock response**

**Table 5.**
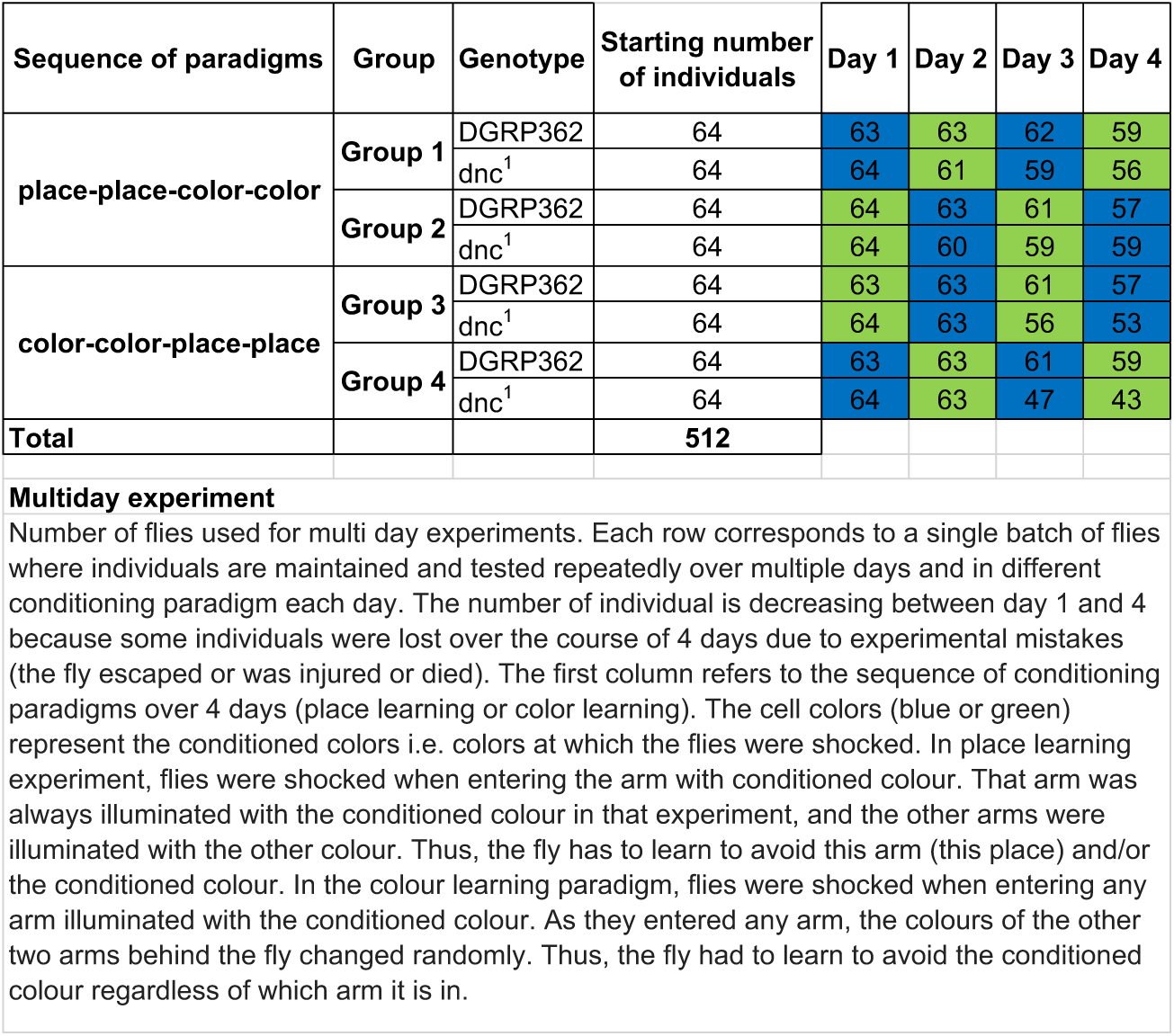
Distribution of individuals - multiday experiment.

**Table 6.**
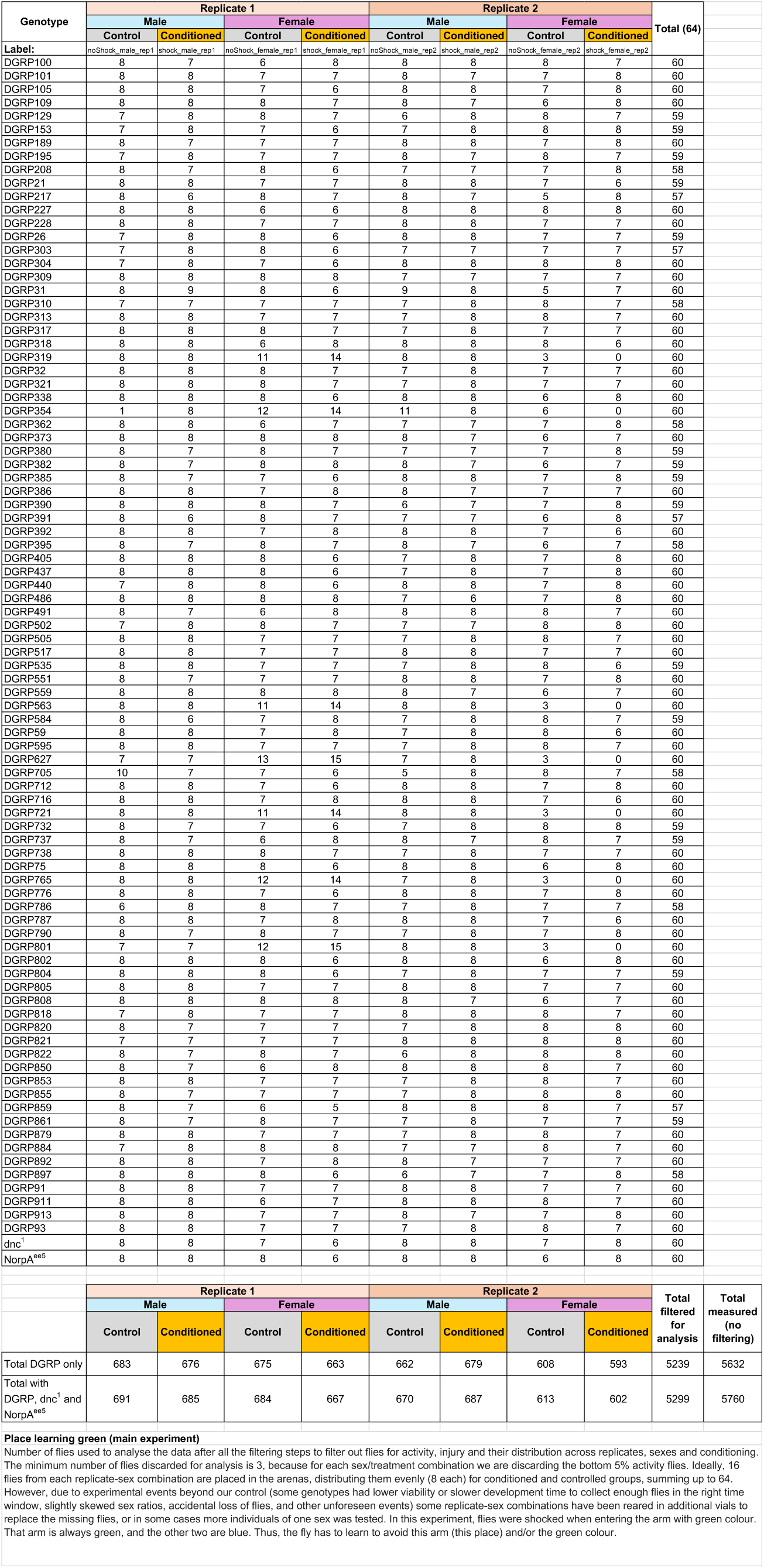
Distribution of individuals - place learning green, main experiment.

**Table 7.**
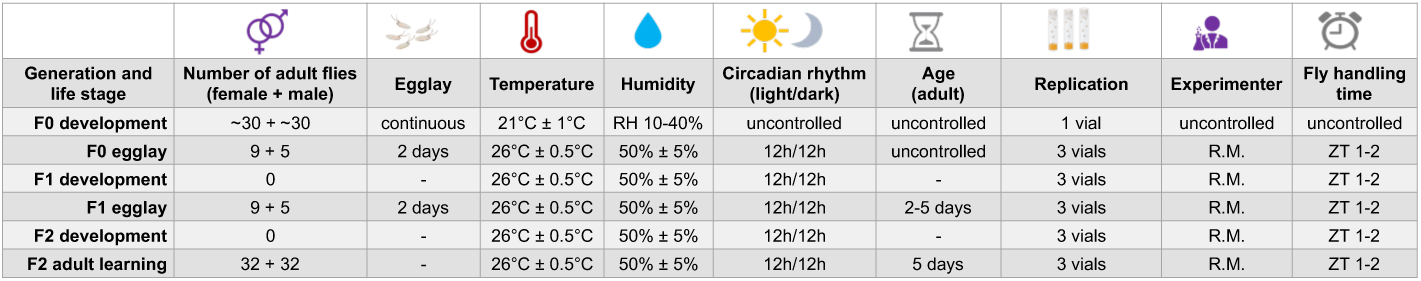
Fly rearing conditions.

**Table 8.**
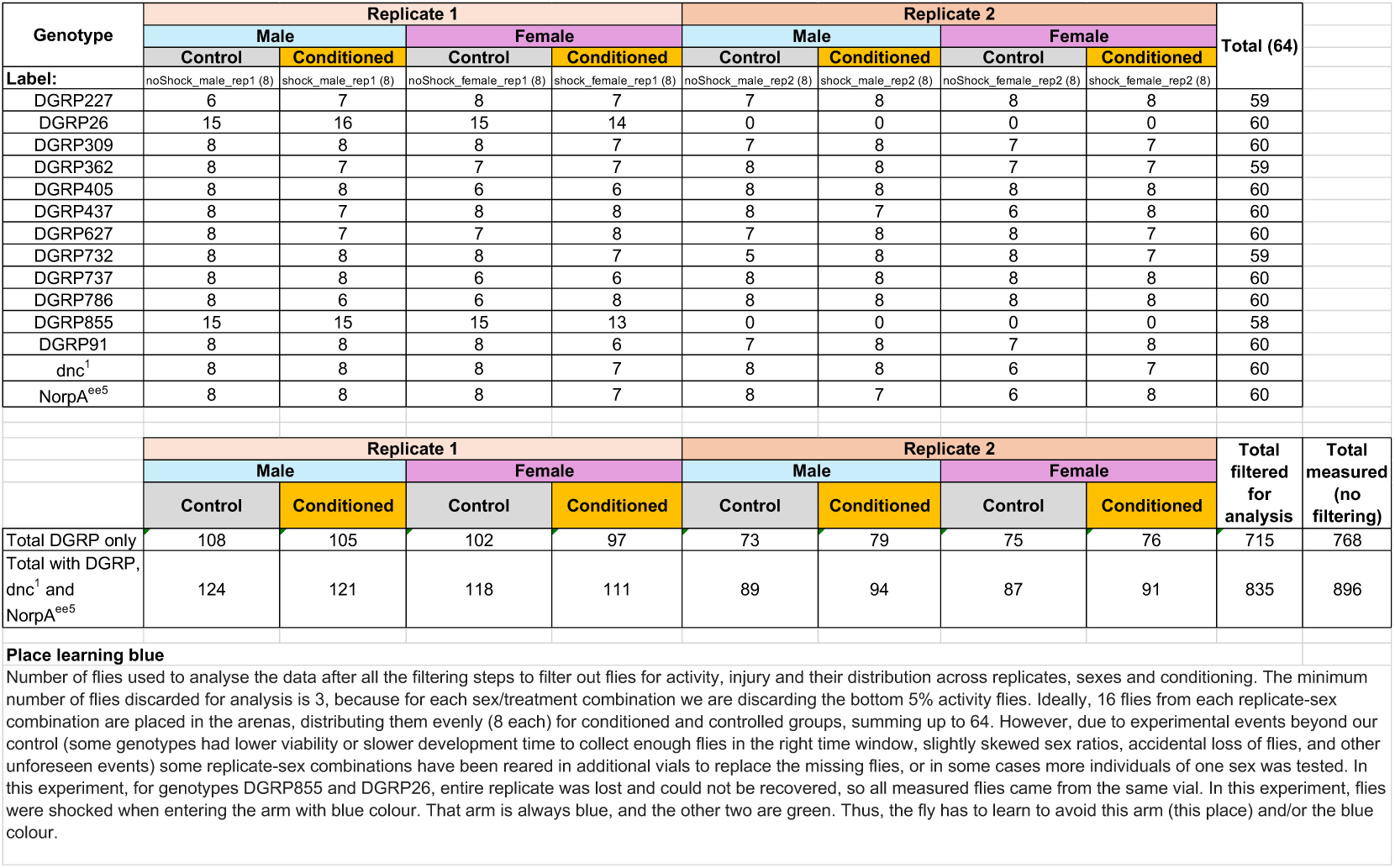
Distribution of individuals - place learning blue, main experiment.

As expected, we found that, on average, conditioned flies spent significantly less time on the colour associated with the shock (23.3%), compared to the control flies (29.9%; N = 2611 and N = 2628, respectively after filtering; T-test p *<* 2.2e^-16^; Figure 2A, Figure 2 - figure supplement 2A, Table 9). This change was lacking in learning- and vision impaired genotypes (Figures 1F, 1G, Figure 1 - figure supplement 1B, Figure 1 - figure supplement 1D, Figure 1 - figure supplement 1E). Given that the memory mutant *dnc^1^* is known to be able to sense shock ***Dudai et al. (1976)***, the observed lack of change in task performance pointed to impaired learning. Oppositely, the majority of wild-type genotypes (87.5%) showed change in average task performance towards avoidance of the shocked colour (Figure 1G; ANOVA p *<* 2.2e^-16^, Sum Sq. = 3.143; Table 9). This change was consistent across sexes, time, and across paradigms (Figure 1 - figure supplement 1C; Table 10).

**Table 9.** Task performance model: green place learning.

| Type III Analysis of Variance Table with Satterthwaite's method |  |  |  |  |  |  |  |
| --- | --- | --- | --- | --- | --- | --- | --- |
|  | Sum Sq | Mean Sq | NumDF | DenDF | F value | Pr(>F) |  |
| genotype | 3.14298 | 0.03613 | 87 | 4901.1 | 3.3359 | < 2.2e-16 | *** |
| treatment | 0.93192 | 0.93192 | 1 | 1808.9 | 86.0524 | < 2.2e-16 | *** |
| shock_sensitivity | 0.05228 | 0.05228 | 1 | 4933.3 | 4.8276 | 0.028053 | * |
| sex | 0.00417 | 0.00417 | 1 | 78.4 | 0.3847 | 0.536921 |  |
| developmental_noise | 0.06312 | 0.06312 | 1 | 4912.5 | 5.8282 | 0.015808 | * |
| starting_arm_code | 0.13062 | 0.04354 | 3 | 4955 | 4.0204 | 0.007221 | ** |
| genotype:treatment | 3.08198 | 0.03543 | 87 | 4873.7 | 3.2711 | < 2.2e-16 | *** |
| shock_sensitivity:sex | 0.02418 | 0.02418 | 1 | 4942.4 | 2.2325 | 0.135201 |  |
| treatment:starting_arm_code | 0.00376 | 0.00125 | 3 | 4955.8 | 0.1158 | 0.950898 |  |

**Table 10.**
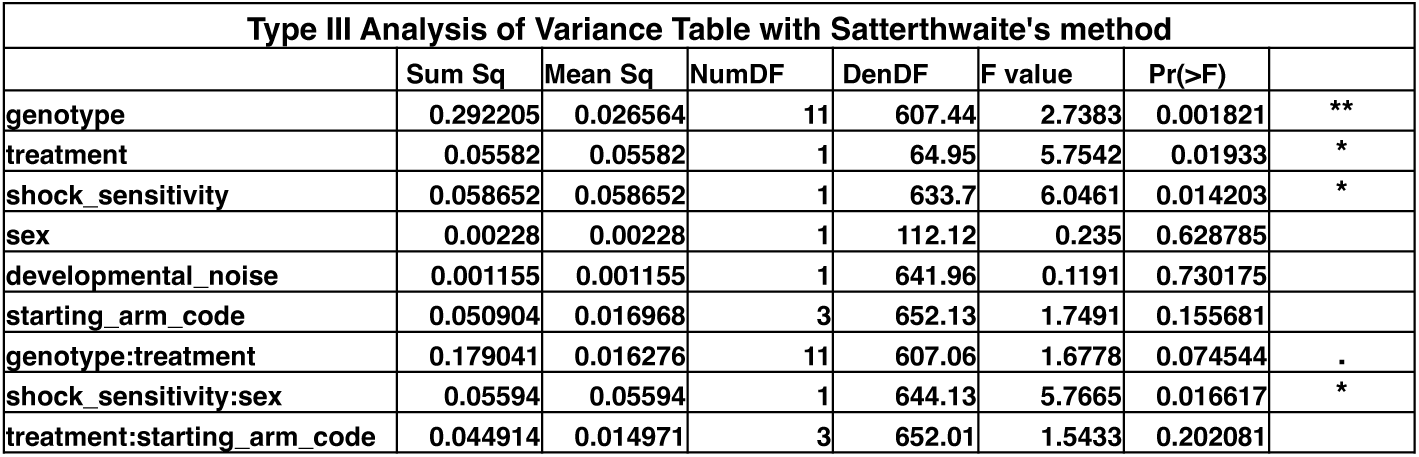
Task performance model: blue place learning.

Most other task-relevant behaviours were also significantly different between control and conditioned flies (Figure 1 - figure supplement 2). These differences were consistent over the time course of the assay, across learning paradigms, replicates and across sexes (ANOVA p = 0.537, Sum Sq. = 0.004, Figure 1 - figure supplement 2, Table 10-11).

**Table 11.**
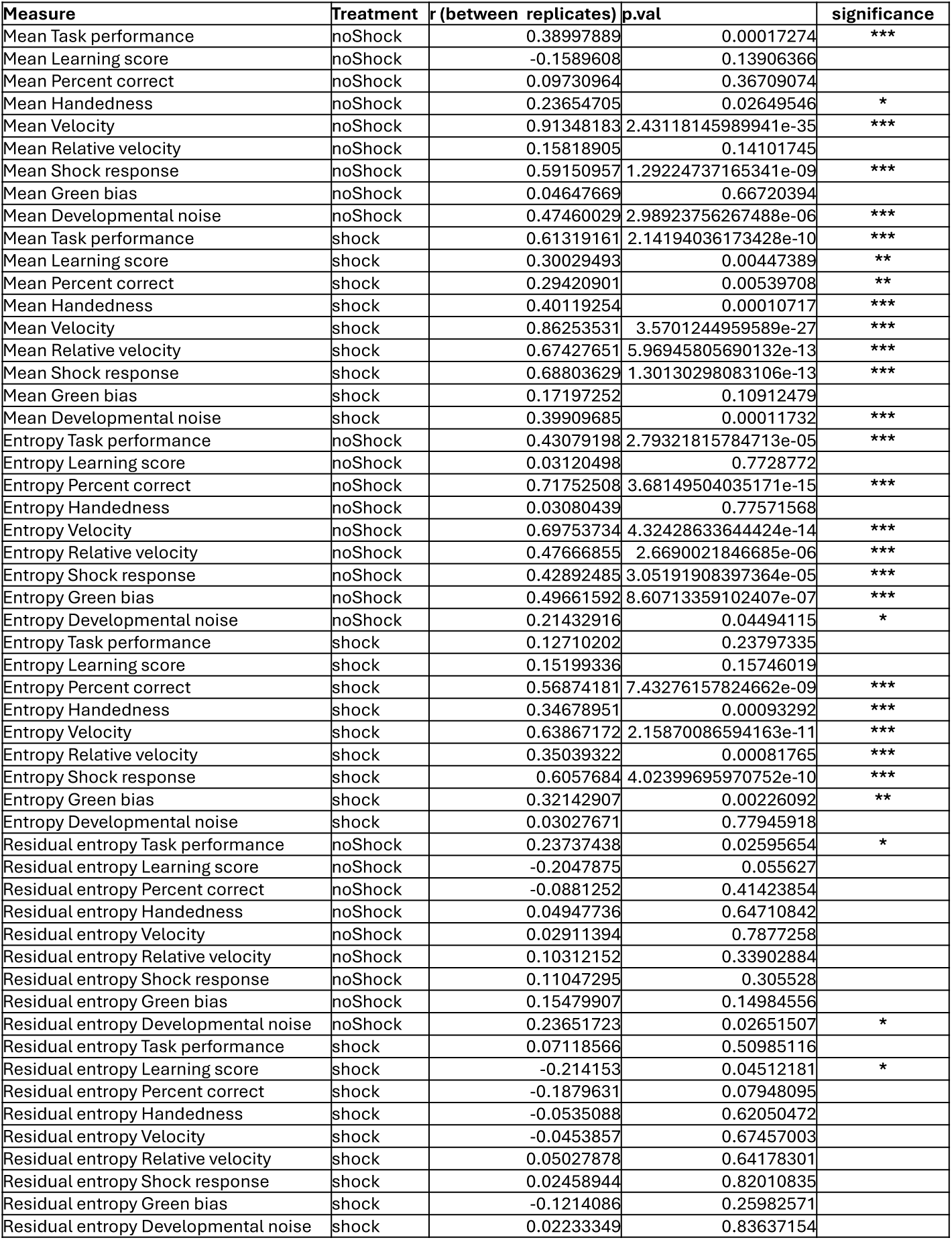
Pearson product-moment correlation and significance between mean and individuality measures of behaviour between replicates (N = 32 flies per genotype per replicate (16 conditioned, 16 control) for 88 genotypes.

| Measure | Treatment | r (between replicates) | p.val | significance |
| --- | --- | --- | --- | --- |
| Mean Task performance | noShock | 0.38997889 | 0.00017274 | *** |
| Mean Learning score | noShock | -0.1589608 | 0.13906366 |  |
| Mean Percent correct | noShock | 0.09730964 | 0.36709074 |  |
| Mean Handedness | noShock | 0.23654705 | 0.02649546 | * |
| Mean Velocity | noShock | 0.91348183 | 2.43118145989941e-35 | *** |
| Mean Relative velocity | noShock | 0.15818905 | 0.14101745 |  |
| Mean Shock response | noShock | 0.59150957 | 1.29224737165341e-09 | *** |
| Mean Green bias | noShock | 0.04647669 | 0.66720394 |  |
| Mean Developmental noise | noShock | 0.47460029 | 2.98923756267488e-06 | *** |
| Mean Task performance | shock | 0.61319161 | 2.14194036173428e-10 | *** |
| Mean Learning score | shock | 0.30029493 | 0.00447389 | ** |
| Mean Percent correct | shock | 0.29420901 | 0.00539708 | ** |
| Mean Handedness | shock | 0.40119254 | 0.00010717 | *** |
| Mean Velocity | shock | 0.86253531 | 3.5701244959589e-27 | *** |
| Mean Relative velocity | shock | 0.67427651 | 5.96945805690132e-13 | *** |
| Mean Shock response | shock | 0.68803629 | 1.30130298083106e-13 | *** |
| Mean Green bias | shock | 0.17197252 | 0.10912479 |  |
| Mean Developmental noise | shock | 0.39909685 | 0.00011732 | *** |
| Entropy Task performance | noShock | 0.43079198 | 2.79321815784713e-05 | *** |
| Entropy Learning score | noShock | 0.03120498 | 0.7728772 |  |
| Entropy Percent correct | noShock | 0.71752508 | 3.68149504035171e-15 | *** |
| Entropy Handedness | noShock | 0.03080439 | 0.77571568 |  |
| Entropy Velocity | noShock | 0.69753734 | 4.32428633644424e-14 | *** |
| Entropy Relative velocity | noShock | 0.47666855 | 2.6690021846685e-06 | *** |
| Entropy Shock response | noShock | 0.42892485 | 3.05191908397364e-05 | *** |
| Entropy Green bias | noShock | 0.49661592 | 8.60713359102407e-07 | *** |
| Entropy Developmental noise | noShock | 0.21432916 | 0.04494115 | * |
| Entropy Task performance | shock | 0.12710202 | 0.23797335 |  |
| Entropy Learning score | shock | 0.15199336 | 0.15746019 |  |
| Entropy Percent correct | shock | 0.56874181 | 7.43276157824662e-09 | *** |
| Entropy Handedness | shock | 0.34678951 | 0.00093292 | *** |
| Entropy Velocity | shock | 0.63867172 | 2.15870086594163e-11 | *** |
| Entropy Relative velocity | shock | 0.35039322 | 0.00081765 | *** |
| Entropy Shock response | shock | 0.6057684 | 4.02399695970752e-10 | *** |
| Entropy Green bias | shock | 0.32142907 | 0.00226092 | ** |
| Entropy Developmental noise | shock | 0.03027671 | 0.77945918 |  |
| Residual entropy Task performance | noShock | 0.23737438 | 0.02595654 | * |
| Residual entropy Learning score | noShock | -0.2047875 | 0.055627 |  |
| Residual entropy Percent correct | noShock | -0.0881252 | 0.41423854 |  |
| Residual entropy Handedness | noShock | 0.04947736 | 0.64710842 |  |
| Residual entropy Velocity | noShock | 0.02911394 | 0.7877258 |  |
| Residual entropy Relative velocity | noShock | 0.10312152 | 0.33902884 |  |
| Residual entropy Shock response | noShock | 0.11047295 | 0.305528 |  |
| Residual entropy Green bias | noShock | 0.15479907 | 0.14984556 |  |
| Residual entropy Developmental noise | noShock | 0.23651723 | 0.02651507 | * |
| Residual entropy Task performance | shock | 0.07118566 | 0.50985116 |  |
| Residual entropy Learning score | shock | -0.214153 | 0.04512181 | * |
| Residual entropy Percent correct | shock | -0.1879631 | 0.07948095 |  |
| Residual entropy Handedness | shock | -0.0535088 | 0.62050472 |  |
| Residual entropy Velocity | shock | -0.0453857 | 0.67457003 |  |
| Residual entropy Relative velocity | shock | 0.05027878 | 0.64178301 |  |
| Residual entropy Shock response | shock | 0.02458944 | 0.82010835 |  |
| Residual entropy Green bias | shock | -0.1214086 | 0.25982571 |  |
| Residual entropy Developmental noise | shock | 0.02233349 | 0.83637154 |  |

Interestingly, we found an extensive variability in expressed individual behaviour within the wild-type genotypes. Distributions of individual task performance were broader in some genotypes than in others (Figures 1E, 1F, 2A; Figure 1 - figure supplement 2A, Figure 2 - figure supplement 1A). This variability appeared more prominent in conditioned wild-type flies. Conversely, we also observed smaller inter-individual variability in task performance in the conditioned learning mutants *dnc^1^* (Figure 1 - figure supplement 2A).

Curiously, not only the means and variances, but also the shapes of the distributions of individual behaviours varied reproducibly between genotypes, particularly in the conditioned group (Figure 2B; Figure 2 - figure supplement 1A). Unlike control flies, the individual behaviour of conditioned flies often occupied non-Gaussian distributions (Figure 2B, Figure 2C; Figure 2 - figure supplement 1A). We reasoned that the process of conditioning, and specifically learning, may be inducing this additional expression individuality within genotype.

However, one obvious difference in the environment between conditioned and control flies is the presence of the shock stimulus which could be perceived individually, thus affecting the shape of the distribution. We tested whether individual sensitivity to shock could account for the observed changes in individual task performances in the two groups. We defined *shock response* as the difference between the velocity of an individual fly measured during the three minutes of bias testing prior to the start of the task and its velocity in the first quarter of the task, when the shock becomes implemented in the conditioned group. This way, instead of using yoked shock response measurement, we preserve within-individual measurement of shock response. Shock response in individual flies did not correlate with their task performance, whether it was measured at the individual (Figure 2 - figure supplement 3C) or the genotype level (Figure 2 - figure supplement 4A). It was, however, expectedly higher in the conditioned flies that actually experienced shock (mean *shock response*_control_ = 0.294 mm/s, mean *shock response*_conditioned_ = 0.451 mm/s, T-test p *<* 2.2e^-16^, n = 5239, Figure 2 - figure supplement 4B, Figure 2 - figure supplement 4C). There was also no correlation between shock response and other task relevant behaviours in the conditioned group (Figure 2 - figure supplement 3, Figure 2 - figure supplement 4A). Therefore, shock response alone is unlikely to account for the change in the distribution of individual task performance. This leaves learning as the more likely explanation for the observed variation in the shape of the distributions.

### Learning exposes residual individuality

Because the shapes of the distributions were not always normal, we could not use the classical measurements of expressed individuality, namely the genotypes’ mean and variance, to compare between groups. Instead, to compare the differences in expressed individuality between any group of flies we use Hellinger distance which captures the differences in shapes of distributions. This way we need not assume Gaussian probability distributions of individual behaviour. Further, we introduce a measure of *differential entropy* (H) to capture the extent of expressed individuality within a group of flies (see Methods for details).

The entropy of task performance was significantly higher in conditioned flies (Figure 2 - figure supplement 5) but correlated weakly, though significantly, between conditioned and control flies across genotypes (Pearson’s product-moment correlation between H_control_ and H_conditioned_ r = 0.232, p = 0.03, n= 88 genotypes, Figure 2 - figure supplement 4A). This indicated that a significant portion of the observed variation in individual behaviour within genotype was sourced from life history processes that preceded the splitting of flies to control and condition group - before the behaviour was measured. On the other hand, the difference in distributions between replicates of the same genotype was equally low in both conditioned and control flies (Mean *Hellinger*_replicates_ = 0.06, Mean *Hellinger*_replicates_ = 0.05, respectively, T-test p = 0.256, n = 88, Figure 2 - figure supplement 4C). This confirmed that the controlled environmental conditions were matched across vials. The observed individual variability is thus primarily sourced from stochastic processes that occurred during development and life prior to the behaviour assay.

The entropy of task performance was strongly dependent on the genotype (Resampling test, parametric p *<* 2.2e^-16^ resampled p = 1e^-06^, Figure 2 - figure supplement 4B). Yet, the extent of expressed individuality (entropy) in spontaneous behaviour was not translated linearly to individu-ality in conditioned behaviour (Figure 2B, Figure 2 - figure supplement 1A, Figure 2 - figure supplement 4A). In other words, the extent of expressed individuality of a genotype differed significantly between innate colour preference and learned colour preference. Higher entropy can reflect a difference in interaction of genotypes with conditioned compared to control task environment; a simple change in the distribution’s mean and variance could result in higher entropy. However, the evident changes in higher-order features of the distributions such as their shape, indicated an additional reorganization of individual behavioural expression between control and conditioned flies.

To capture and quantify this additional expression of individuality we introduced a measure of *“residual individuality”* (H_resid_). H_resid_ is calculated as Shannon entropy of a probability distribution after scaling the distribution within its own range (Figure 2D, Figure 2 - figure supplement 6). By scaling the distributions within their range, this measure nullifies the variability induced by factors shared across all groups of flies, including stochasticity in rearing environments, genotype, and their interaction, which are normally read out from measures of central tendency and dispersion (Figure 2 - figure supplement 6). With such scaling we could measure the residual changes in the shapes of the distribution, independent of their ranges and means.

Besides the measure of residual individuality within a distribution, we used three different mea-sures of shape divergence between distributions (Δ H_resid_) to dissect the source of the residual individuality. *Learning divergence* (Δ *H_resid_ Learning*) quantifies the change in expressed residual individuality of a genotype that arises due to learning (H_resid_ conditioned vs. H_resid_ control; Fig-ure 2D). *Experimental divergence*, (Δ *H_resid_ Experimental*) reflects the change in H_resid_ that may have been caused by some experimental factor affecting flies individually but systematically differently across vials (H_resid_ replicate 1 vs. H_resid_ replicate 2). *Genetic divergence*, (Δ *H_resid_ Genetic*) captures the genetic contribution to the expressed residual individuality observed during the behaviour assay (H_resid_ genotype A vs. H_resid_ genotype B) (see Methods for details on all three measures).

We found no difference in *experimental divergence* between conditioned and control groups (Δ *H_resid_ Experimental*_control_ = 2.08, Δ *H_resid_ Experimental*_conditioned_ = 1.85, T-test p = 0.206, n = 88, Figure 2I). Technical conditions of the experiment thus did not cause significant changes in the shape of the distributions of conditioned flies. On the other hand, *Genetic divergence* revealed that genotype contributed to the shapes of the distributions of task performance. The shapes were significantly more divergent across genotypes in the conditioned flies (Δ *H_resid_ Genetic*_control_ = 0.70, Δ *H_resid_ Genetic*_conditioned_ = 0.83, T-test p *<* 2.2e^-16^, n = 88 genotypes; Figure 2H). This was also true for the *learning divergence* (Δ *H_resid_ Learning*), which differed significantly across genotypes (Resampling test, parametric p *<* 2.2e^-16^, resampled p = 1e^-06^; Figure 2E).

These results were unexpected; in a strictly controlled environment shared between individu-als, genetic differences are expected to be the only deterministic variables that can reproducibly affect expressed individuality. More strictly, in isogenic populations in a shared environment, the effect of a genotype on individuality is assumed to be absent except for its effect on phenotype’s variance. This is primarily a result of the genotype-specific buffering of stochastic processes during the development. In both cases, individuality is expressed and read out only through changes in the mean and variance of a distribution, but not its shape ***Ayroles et al. (2015)***; ***Linneweber et al. (2020)***. However here, the mean and variance was accounted for, and yet we still find systematic differences in expressed residual individuality in task performance. We found this to be true for other behaviours as well (Figure 2 - figure supplement 1E-I, Figure 2 - figure supplement 7). We could not explain this variation with residual heterozygosity (Figure 2 - figure supplement 4A) or response to shock (either individual or genotype-specific; Figure 2 - figure supplement 3, Figure 2 - figure supplement 4A). However, we found that *learning divergence* also evolved within each genotype with every made decision (Figure 2F). This revealed that the dynamics of decision-making may help pinpoint the source of this additional individuality.

### Learning and bias amplify the butterfly effect of individual experience

Despite the efforts to fully control and parallelize experience, perfect control over freely behaving biological agents is by definition not possible. In non-uniform environments, momentary experience of a freely behaving individual will depend on its previous decisions. As a result, the sequence of decisions made in a natural environment is too diverse to be tractable. The Y-maze arena, however, minimizes the diversity of unique experiences and binarizes the decisions a fly can make. This allowed us to track and enumerate decision-making (Figure 3B, 3C).

**Figure 3.**
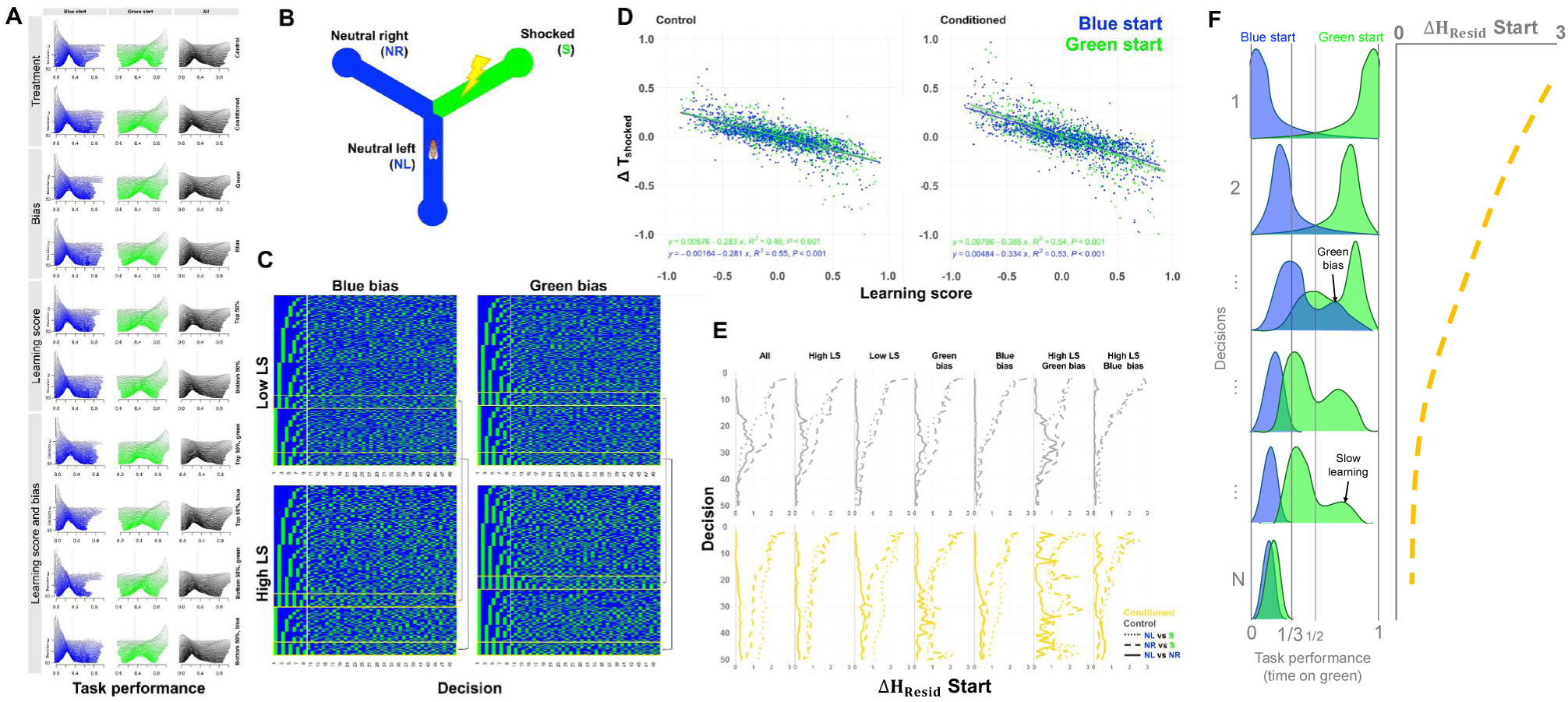
Divergence in individual task performance depends on learning through experience. (A) Task performance distributions over the first 50 decisions is dependent on starting colour despite their initial colour bias or their final learning score (LS). The distributions of flies starting at the left and right blue arm are overlaid. Only flies that made at least 50 decisions are included. Distributions regardless of starting position are shown in black. (B) Schematic of the Y-maze and starting colours. For the conditioned flies, the green coloured arm (S) is associated with the shock, while for control flies it is not. (C) Alignment of 50 decisions made by conditioned flies (rows) with different learning score and colour bias combinations. Decisions are sequentially sorted. The yellow rectangles highlight flies with most persistent behaviour of alternating choices between colours. Black lines indicate flies that share the same first decision for easier comparison across groups. (D) Change in individual task performance depends more strongly on initial conditions for flies that learn. Each point is an individual fly. (E) Evolving change in residual individuality between flies starting at different arms of the Y-maze, that made at least 50 decisions (H_resid_ *Start*). Residual individuality is sensitive to initial experience and exacerbated by opposing forces of learning and unfavourable bias. (F) Schematic showing how the shape of the distributions with every made decision may change for flies that started the assay at different colours and how this may translate to ΔH_resid_ *Start*. Flies that learn will more quickly shift the distribution to the right. On the other hand, green-biased flies will push the distribution to the right. **Figure 3—figure supplement 1. Effect of starting position, innate colour bias and learning on task performance** **Figure 3—figure supplement 2. Distribution of T_shocked_ of the conditioned (shocked) flies over different periods of the assay.** **Figure 3—figure supplement 3. Distribution of T_shocked_ of the control (not shocked) flies over different periods of the assay.**

We aligned discrete decision sequences across thousands of individuals to ask whether mo-mentary experience drives behavioural divergence. This uncovered that even minimal, single-moment differences in individual experience, such as the initial conditions of the first decision, can strongly shape learning trajectories. Because flies move freely, and the task onset occurs at an arbitrary but common timepoint, each individual can begin the task from any arm. A fly that finds itself on the green arm will immediately experience shock in the very first second, at the start of the task. In contrast, a fly that starts in the blue arm will experience its first shock only after making its first incorrect decision. Thereafter, the consequences of subsequent decisions are identical across individuals. This bears an effect on individual task performance dynamics and amplifies individuality over time (Figure 3A, Figure 3D, Figure 3E, Figure 3F; Figure 3 - figure supplement 1D).

Early stochastic differences in the first decision provide a plausible mechanism by which residual individuality diverges between control and conditioned flies and across genotypes, despite the otherwise shared environment. Although the initial arm choice appeared random at the individual level, starting positions were normally distributed across genetically identical flies, with genotype-dependent means (Figure 3 - figure supplement 1A, Figure 3 - figure supplement 1B, Figure 3 - figure supplement 2, Figure 3 - figure supplement 3). Each decision can therefore be conceptualized as a single draw from an individual-specific probability distribution of colour preference, partially determined by genotype. With each subsequent decision, this distribution is resampled and, if learning occurs, updated as a function of experience. Consistent with this framework, we found that learning progressively outweighed innate starting bias as the determinant of individual task performance (Figure 3F, Figure 3 - figure supplement 1E-F, Figure 3 - figure supplement 2, Figure 3 - figure supplement 3).

Among conditioned flies sharing the same initial biases, an early stochastic event could trigger an earlier or qualitatively different learning trajectory in one individual but not another. In control flies, which do not learn, decision rules and their consequences on task performance would be insensitive to initial conditions. Accordingly, in control flies time-averaged distribution of task performance converged to the same shape despite differences in first decisions. In conditioned flies, residual individuality originating from the first decision persisted across many subsequent choices (Figure 3E). We found that the distributions of individual behaviour were broader and their shapes changed substantially over the course of the experiment for the conditioned flies, and not for the control flies Figure 3 - figure supplement 2, Figure 3 - figure supplement 3).

To further characterize the potential causes of this effect, we grouped flies by initial colour bias and by final learning score. We then measured change in residual individuality (Δ *H_resid_ Start*) in task-performance between individuals starting the task from different arms. Divergence was maximized at the first decision (Figure 3E, Figure 3F). In control flies, this divergence rapidly collapsed despite distinct initial conditions or initial biases. In conditioned flies, divergence persisted and was further amplified in individuals with high learning scores and an unfavourable initial biases (Figure 3E). Alignment of decision sequences further revealed an early alternating-choice pattern in conditioned flies, particularly pronounced in individuals with an initially unfavourable bias who ultimately achieved high learning scores (Figure 3C). This suggested that the clash between learning and innate bias likely contributes to residual individuality. These dynamics indicate that the learning experience of the first decision can initiate a butterfly effect: because learning is non-ergodic, the effects of early stochastic differences accumulate over time, progressively diversifying experience and amplifying individuality within a genotype.

### Learning diversifies individual behaviours

While we do not track behaviour throughout the entire life of a fly, the design of our experiment allows us to dissect the effects of life experience prior to the task from the effects of momentary experience during the task. We asked how extensive is the effect of the entire life’s experience on individual momentary behaviour, compared to the effect of genetics or learning during the task? To quantify the relative contributions of different sources of individuality, we turned to modelling of observed behaviours. To test how well our models capture and represent the observed behaviour we used them to simulate behaving flies and then compared their simulated behaviour to the behaviour of observed, real flies. Simulated flies incorporated a range of behavioural parameters (Table 12). Based on them, the simulated flies decided how long they want to stay in the current arm of a simulated Y-maze, and whether they want to turn left or right when leaving the current arm. The left-right turns were sampled from a Bernoulli distribution with a time-dependent rate. The decision to leave the arm was made by sampling the escape duration (time to leave one arm) from a log-normal distribution with a time-dependent mean and stationary variance. The mean of their escape duration distribution depended on the colour of the arm and the presence of shock at the time point *𝑡*. Importantly, the mean escape duration and the rate of turning left or right could change with a reinforcement learning rule, which depends on the presence of shock.

**Table 12.** Theoretical ranges of measured behaviours.

| Behavior | low | high | unit |
| --- | --- | --- | --- |
| Task performance | 0 | 1 | - |
| Learning score | -1 | 1 | - |
| Percent correct decisions | 0 | 1 | - |
| Relative velocity | -8 | 8 | mm/s |
| Handedness | 0 | 1 | - |
| Velocity | 0 | 8 | mm/s |
| Green bias | 0 | 1 | - |
| Developmental noise | 0 | 1 | - |
| Shock response | 0 | 8 | mm/s |
| $\Delta$ Task performance | -1 | 1 | - |
| $\Delta$ Percent correct decisions | -1 | 1 | - |
| $\Delta$ Handedness | -1 | 1 | - |
| $\Delta$ Relative velocity | -8 | 8 | mm/s |

To assess the effects of genetics, learning, and individual life experience, we systematically exclude one or none of them in four different behaviour models. First, all flies of the same genotype have identical parameters, but different genotypes have different parameters (*no individuality* model), thereby omitting the effect of any individual experience obtained prior to expressing a behaviour in the task. Second, the *𝑖*-th parameter of an individual simulated fly is sampled from a normal distribution with genotype-dependent mean (*𝑚_i_*^(^*^x^*^)^) and standard deviation (*𝑠_i_*^(^*^x^*^)^). These simulated flies have individuality in the sense that their initial behavioural policies differ from one another, because no two flies have the exact same parameters (*individuality* model). In other words, simulated flies start the task as if in addition to their genetics and learning, their life experience prior to the task (including stochastic processes during development) had shaped their individual behaviour. Third, parameters are sampled as in the *individuality* model, but the learning rate is set to zero, such that the presence of shock or colour induce no lasting change on their behaviour beyond the immediate effect of changing the escape duration, simulating only the shock response (*no learning* model). Fourth, parameters are sampled per individual, but there is no genotype dependence of the means and standard deviations of the parameter distributions. This way the effect of genetics is omitted, including its effect on developmental variability (*no genetics* model).

Best fit was found for *individuality model*, followed by *no learning* and *no genetics* model. *No individuality model* failed to recapitulate the observed behavioural individuality (Figure 4A-B). Particularly striking was the mismatch in the distribution of the number of decisions made by flies simulated without individuality compared to the actually measured distributions of the number of decisions made by real flies (Figure 4 - figure supplement 1). Compared to whole-life experience or genetics, the effect of momentary learning on individual behaviour was much smaller, and for some genotypes seemed irrelevant (Figure 4A). However, we found that the model without learning could not capture the temporal dynamics of many observed behaviours, highlighting its significance despite apparently modest effect size (Figure 4C). Interestingly, the excess of observed non-Gaussian distributions of individual behaviour in conditioned flies was recreated only if the simulated flies could learn, despite the effect of shock on activity being simulated in no learning model, and despite all parameters of the model being drawn from normal distributions (Figure 2C). Lastly, repeated simulations revealed that the genotype distribution of simulated task performance matches the observed behaviours better for the model with learning (Figure 4D, Figure 4 - figure supplement 1-2). This reaffirmed that residual individuality most plausibly emerges from the very process of learning.

**Figure 4.**
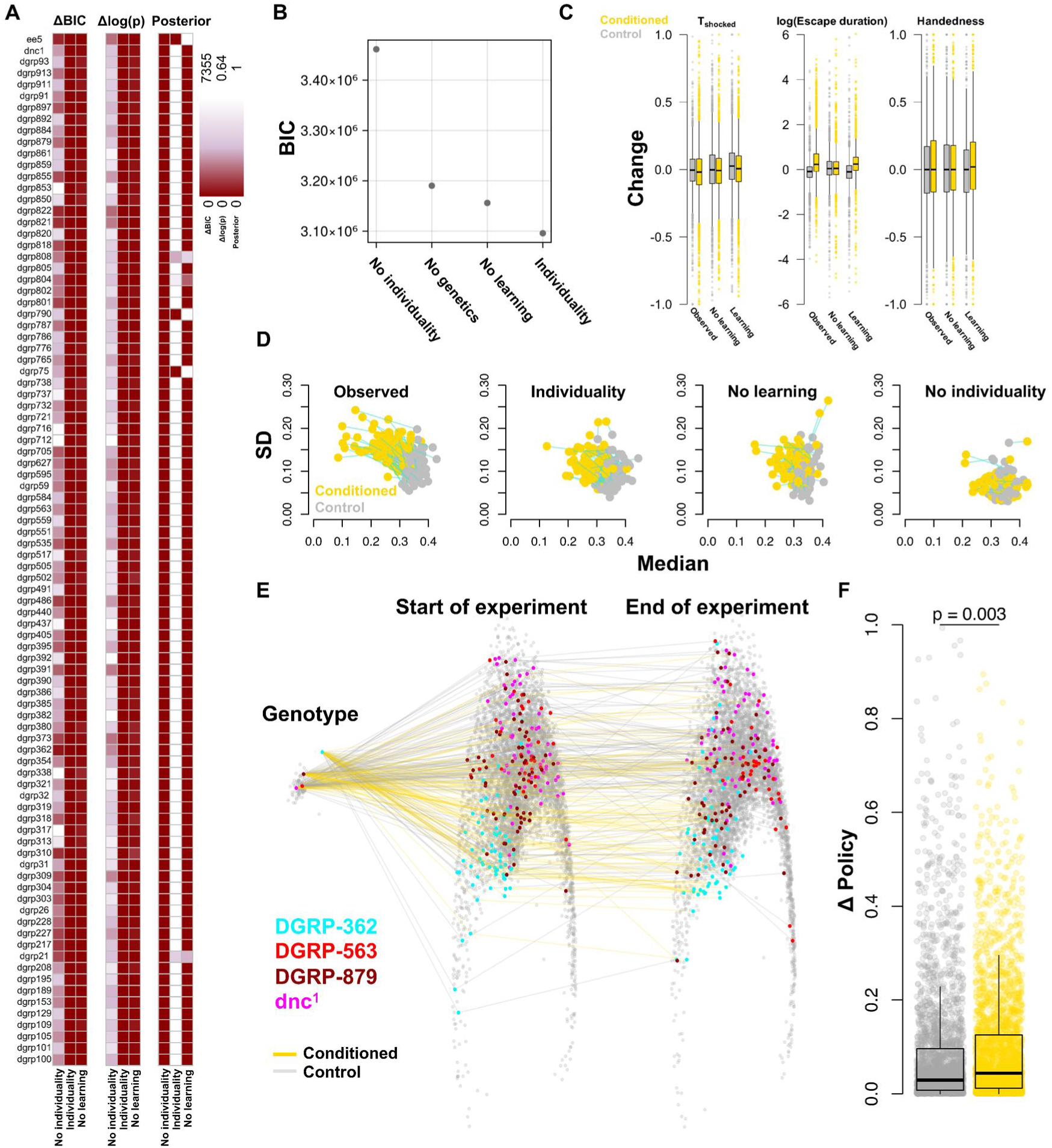
Experience, genetics and learning shape individuality. (A) Variation in behaviour observed across genotypes is best explained by a model that includes genetically controlled but individually random prior life experience, and learning (*individuality*). (B) High Bayesian information criterion for models omitting variability in behaviour prior to the task, genetics, and learning demonstrate that they are necessary for close *in silico* recapitulation of observed individuality. (C) Change in individual behaviours during conditioning can be attributed to the effect of learning. Boxplots summarize the behaviours of observed, real flies and of flies simulated with and without learning. (D) Flies simulated under the assumption of genetic control over variability in life experience and with learning during task can faithfully reproduce observed mean task performance and individual variation in behaviour of real flies. Points are genotypes. (E) Genetic effect on behavioural policy projects to the entire individual behaviour space. Mean genotype policies and overlapping individual policies measured at the start and at the end of the experiment are projected on a multidimensional scaling plot. Points show genotypes and individuals. Highlighted points correspond to individuals from an outlier genotype (turquoise), two average genotypes (red and dark red) and a memory-deficient genotype (magenta). (F) Change from initial behavioural policy from the start until the end of the task changes more in flies that learn. Points are individuals. **Figure 4—figure supplement 1. Summary statistics for observed and *in silico* behaviours.** **Figure 4—figure supplement 2. Quantification of posterior predictive checks.** **Figure 4—figure supplement 3. Influence of behavioural policy on task performance.**

The fitted models allowed us to estimate how the momentary task-relevant behaviour of a given fly differs from other flies, or how it compares to other moments in time. We found that the individual behaviour had changed significantly during the experiment (Figure 4E-F) and that this change was significantly bigger in the flies that learned (Figure 4F, Figure 4 - figure supplement 3). A visualization of these behavioural differences with multidimensional scaling revealed that the behaviour of individual flies does not cluster according to genotype throughout the experiment (Figure 4E). This lack of clustering was also confirmed with a classification analysis: using a long-short term memory (LSTM) classifier applied to the behavioural data of individual flies, we could predict the genotype in only 3.4% of the cases (Chance level : 1.1%, 4-fold cross-validation). Although our models confirmed that genetic variation is essential for generating the observed behavioural diversity, they demonstrate it determines an almost stochastic fraction of any given individual’s behaviour.

Altogether, these results decisively demonstrate that learning is not just an emergent but a fundamental driver of behavioural individuality.

## Discussion

With genetically identical flies raised and tested under identical conditions, we showed that even short learning experiences contribute to behavioural diversification. Our findings reaffirm the well-established importance of developmental environment and early life genotype × environment interactions in the emergence of individuality ***Linneweber et al. (2020)***; Thomas F. ***Mathejczyk et al. (2023)***; ***Maloney et al. (2024)***; ***Laskowski et al. (2022)***. However, we further demonstrate that individual, momentary learning experience substantially extends behavioural variability beyond the the effects of past events experienced prior to the behaviour. Importantly, we identify learning as a distinct and critical form of genotype × environment interaction that is intrinsically individual because in practice, it cannot be shared between individuals. It is therefore fundamental to understanding behavioural divergence between individuals. We reveal this aspect of learning by measuring residual individuality, the persistent inter-individual variability within a genotype that is leftover after accounting for variation arising from past life experience. This finding challenges the common assumption that future behaviour could, in principle, be reliably predicted from genetics if the past environment was perfectly controlled. The process of learning introduces a currently intractable amount of stochasticity to individual behaviour.

We find that observed behavioural individuality cannot be understood or recreated *in silico* without acknowledging genetic diversity. And yet we could not predict a genotype based on individual behaviour. This is not a contradiction: in accordance with previous studies, we find that genetics merely bounds behavioural expression across individuals ***Ayroles et al. (2015)***; ***Linneweber et al. (2020)***. While a genotype defines a landscape of possible individual behaviours, it does not specify which region of this landscape an individual will occupy; this is largely determined by the individual’s unique experiences across life. Notably, we found that these behavioural landscapes substantially overlap across genotypes, implying that nearly any measured behavioural phenotype can, in principle, emerge from any genotype, even under shared rearing conditions. This perhaps “inconvenient truth” has long been implicitly recognised by researchers studying learning-dependent behaviour: even under tightly controlled experimental conditions, behavioural variability, particularly in cognitive performance, is inevitable and unpredictable, despite genetic or environmental uniformity. In our study we could directly observe how genotype can bias the mean initial conditions of behaviour (e.g., colour preference determining starting arm). Yet, the trajectory and evolution of individual behaviour still remain substantially driven by differential learning from individually stochastic events that are independent of genotype (such as the task initiation). We saw that the flux between genotype, environmental stochasticity and learning can give rise to behavioural instability, where a differential change in bias through learning drives divergence between individuals. Similar to the effect of initial learning experience that we found, a recent study by ***Rozenfeld and Parnas (2024)*** had found that experiencing classical conditioning can influence the efficiency of future operant learning ability. We speculate that if the effect of classical conditioning as observed by Rozenfeld and Parnas is equivalent to our initial conditions effect, this could point to a potential for “one-shot” learning ability in flies. Understanding these phenomena across environments and genotypes could potentially bring us closer to predictability of learning-dependent behaviours.

We attributed learning to be the source of residual individuality. This is because analyses of individual behaviour before and during the learning task, together with reinforcement-learning simulations, indicated that learning, rather than shock response is the primary driver of this inter-individual behavioural divergence. Nevertheless, it is possible that individual sensitivity to shock could map to individual task performance in a non-linear fashion and contribute to behavioural divergence. Such effect would not be detectable through correlation analyses or simulations. Classically, this distinction between learning and shock response would be tested using a yoked shock-control design. However, at the level of individual flies, such controls become ill-defined: pairing conditioned and control individuals with fully matched individual behavioural profiles is infeasible given the definition of an individual. A random pairing of one individual that triggers the shock through decisions with another individual that simply experiences shock would also easily conflate shock sensitivity with unrelated and unmeasured individual differences. One potential future approach would be to assay shock response during an additional bias phase before the learning task, in which shock is uncoupled from colour. However, as we see in our study, even a single shock experience could still alter subsequent learning dynamics and introduce new variation in initial learning conditions, making this approach as flawed as any other (or even misleading). Because behaviour was tracked over a 20-minute window, it also remains unclear whether the observed behavioural divergence would persist over longer timescales, once learning has stabilized across individuals. For the same reason, genetic predictability may be only transiently reduced in this task, rather than fundamentally absent across all behavioural regimes or timescales. Longer-term tracking and broader sampling of behavioural landscapes will be essential to address these questions in future studies. In the same vein, simultaneous measurement of behaviour across a much larger sample of genetically identical individuals and under even stricter environmental conditions would reveal if our estimates of the extent of residual individuality is over- or underestimated.

What we demonstrate here is that when individuals learn, even seemingly trivial early events such as the context of a first decision can trigger a butterfly effect that persistently shapes future behaviour and future events, through closed feedback loops. These dynamics seem to be self-generated and contained at the level of the individual, underscoring the active role of the agent in constructing its own behavioural trajectory. Because learning is experience-driven, cognitive behaviour is set to be inherently individual. In natural settings it is likely fundamentally unpredictable. To uncover the mechanisms and especially the genetic underpinnings of cognition in autonomous agents, we must move beyond population averages, linear models, and snapshot observations within genotypes or species. This need becomes even more pressing in complex animals such as humans, whose lived experiences vastly exceed the simplicity of a fly living in a vial and choosing between two colours in a Y-maze.

## Methods and Materials

### Y-mazes

behavioural arenas were 3D printed in black Tough PLA (Ultimaker B. V., Utrecht, Netherlands) using desktop Fused Deposition Modeling (FDM) 3D printer (Ultimaker S5, Ultimaker BV, Utrecht, Netherlands), with a z-axis resolution of 0.1 mm, and 100% grid infill pattern. The mazes are in the form of the letter ‘Y’, with equal arm lengths. The arms are connected at the center with an angle of 120° from each other, and length = 10 mm and width = 2 mm. This specific arm width was chosen so that the flies are not able to turn around in the middle of the arm. This way the fly must traverse the whole arm and reach the end, where it is provided with a circular turning area of a diameter = 4 mm. This is done so that the fly, after making a decision, remains in the arm for long enough to associate the electric shock with the colour of the arm. For the same reason, the fly only gets shocked once it has entered an arm with its full body length, and not at the junction of the three arms. The same requirement is implemented for the removal of electric shock - the fly needs to exit the shocked arm but also enter the non-shocked arm with the full body length in order to escape the shock (Figure 1C). The total height of the maze within which the fly is restricted is 1.2 mm. The wall of the maze does not erect straight up to the ceiling, but in a cross-section forms a step at 0.8 mm. This is done so that the fly, once resting on the floor of the maze, is not able to climb up to the ceiling of the maze and avoid getting shocked. The ceiling of the maze is a regular 25 x 75 mm glass slide coated on one side (facing the maze) with dry Polytetrafluoroethylene (PTFE) aerosol (KONTAFLON 85, Kontakt Chemie, CRC Industries Europe BV, Zele, Belgium). The coating makes them hydrophobic, and hence, slippery for the flies to walk on. Each slide covers two Y-mazes horizontally. Eight Y-mazes (4 rows x 2 columns) are 3D printed together, with each Y-Maze restricted within an area of 30 mm x 30 mm, which corresponds to an individual shocking area on the transparent shocking grid (described in the next section). A single plate consisting of 8 Y-mazes is then laid over, and glued onto a transparent shocking grid. Eight such arenas (64 Y-mazes) are placed on top of the LCD for the assay.

### Conditioning platform

Indium tin oxide (ITO) coated (400 nm coating corresponding to 4-6 Ω/sq) glass plates (150 mm x 100 mm x 1 mm), were laser etched to form a grid of 8 (4 rows x 2 columns) isolated 30 mm x 30 mm shock delivery pads for each Y-maze. The shocking pads were patterned in the form of an interdigitated grid, with a central pad connected to the ground, and 8 isolated pads connected to an array of normally off electromagnetic relays. Each isolated pad was etched in an interdigital pattern with 15 digits of 1 mm width on each side (ground pad, and active pad connected to the relay) separated by a distance of 20*𝜇*m between each digit. Holes (diameter 0.8 mm) were laser cut into the glass plate at positions that correspond to the center point of the circular ends of the Y maze arms. The holes are used to deliver CO_2_ to anesthetize flies for fly retrieval at the end of the assay. The designed plates were manufactured at Diamond Coatings Ltd., UK. The 3D printed Y-mazes are assembled with the glass plate using adhesive tape or superglue, making sure the glue does not touch the ITO coating. Each such assembled plate is placed on a 3D-printed holder fixed on top of an LCD screen (LG 25BL56WY, 25”, 1920 x 1200 pixels). The holder can hold 8 glass plates and is fitted with grooves for electrical wiring and place marks for copper tapes (12 mm width). Copper tape is applied on the glass plates to make electrical contact between the ITO pads and the copper tapes on the holder. Custom-made printed circuit boards fitted with 5.1 mm round head gold spring contacts (P/N 5099-D-2.0N-AU-1.3C, PTR Hartmann, Germany) electrically connect the copper tapes to the electromagnetic relays, ultimately connecting it to each isolated shocking area. Electrification of the 8 isolated Y-mazes is controlled by an array of 8 relays, which in turn is controlled by an 8-bit shift register (TI 74HC595). The 64 mazes are controlled by 64 individual relays, which are controlled by 8 shift registers. The registers are controlled by the computer program (testY_LCD.py) via an Arduino Mega 2560 Rev. 3 (ATmega2560) (loaded with Arduino_yMaze_shock_595.ino) in closed loop with the position of the fly inside the maze and the visual stimulus presented to it via the LCD screen. Visual stimuli can be, but are not limited to, colours, colour gradients, patterns or light intensities. The whole system is made to be modular, and the plates holding the Y-maze arenas wireless, so that it is not fixed to the LCD screen. This way it can be moved around for the purpose of fast loading the arenas with flies. When electrified, the floor of each maze delivers a constant 30 V DC through the flies legs. The current is limited to 500 mA distributed between 64 arenas (on average 7.8 mA per fly) and powered using a bench-top power supply (RND 320-KA330-5P, RND Lab, Switzerland). The electronics circuits are powered using a 5 V, 90 W switch mode power supply (LRS-100-5, Mean Well Enterprises Co. Ltd., New Taipei City, Taiwan). This current was chosen through optimization experiments in which the goal was to maintain reproducible shock avoidance in flies across several randomly chosen wild-type genotypes, while the memory mutant *dnc^1^* flies, that usually experience most shocks due to their inability to learn, do not get injured during the experiment.

### Closed-loop control

Each Y-maze is associated with an Augmented Reality University of Cordoba (ArUco) marker ***Garrido-Jurado et al. (2014)***, which is identified by the control software (getBoxes_aruco_64.py) to acquire the position of each maze. This is done both to automate the experiment and to maintain the accuracy in case of an accidental misalignment of the plate within the tolerance of the plate holders when setting up the experiments. The position of each moving fly is captured using a custom background subtraction algorithm (unit_fly.py), and fed into the control algorithm (testY_LCD.py, unitMaze.py). In turn the algorithm changes the visual stimulus presented to each fly via coloured patterns on the LCD screen (LCD.py). The algorithm also controls the electric shock delivery for each fly via an Arduino Mega 2560 Rev. 3 in real time (Arduino_yMaze_shock_595.ino). The platform is illuminated with an array of 12 x 8 infrared (940 nm) LED (Figure 1B). The video is captured at 6 fps using a grayscale industrial USB 3.0 camera without an infrared filter (DMK 38UX253, The Imaging Source, Germany), fitted with an infrared long pass filter (dia. 40.5mm thread, 0.5 mm, 093 IR Black 830, Schneider Kreuznach GMBH, Germany). The filter allows the program to ignore any background illumination changes inside the room that can interfere with the accuracy of the fly detection algorithm. The program is generally robust to light interference caused by movement around the setup, but to minimize any potential light interference, as well as to minimize influencing flies’ behaviour the experimenter left the room during the assay or wore black or dark clothes. Different associative learning protocols can be implemented with this system, whose codes can be found on our GitHub repository https://github.com/jaksiclab/GeneticsOfLearningIndividuality. At the end of each protocol, the program outputs a folder containing comma-separated values (.csv) files reporting the data acquisition times (in nanoseconds starting from the time of acquisition of the first fly position), the coordinates of the fly, the arm in which the fly resides at each moment, whether the arena is electrified or not, the correct, incorrect and total number of choices the fly has made at that point of time. The program also automatically outputs the final task performance scores of the flies at the end of the experiment, the time at which the experiment was performed, sped up infrared video recordings of the arena (.mp4) and setup metadata that needs to be input by the user at the start of the experiment. For further automation, this information can directly be accessed by the control software immediately after an experiment.

### Drosophila stocks and rearing

All genotypes were acquired from the Bloomington Drosophila Stock Centre (BDSC) in the year 2020. 88 Drosophila Genetic Reference Panel (DGRP) lines, norpA^EE5^ (BDSC stock number 5685) mutants, and *dnc^1^* (BDSC stock number 6020) mutants were used in this study. DGRP genotypes are denoted by a number (*nnn*) or a label *DGRP-nnn*, where *nnn* represents the DGRP genotype. For example, in Figure 2D and 2H, genotype was denoted as 208 or DGRP-208, respectively, and both indicate that the genotype used was DGRP-208 line (synonym RAL-208; BDSC stock number 25174). Flies were maintained and raised on cornmeal agar medium: 6.2 g agar-agar, 58.8 g cornmeal, 58.8 g brewer’s yeast, 0.1 L grape juice - Migros M-Classic, 4.9 mL propionic acid, 26.5 mL methylparaben Moldex, 1 L of tap water (Lausanne, Switzerland). Environmental temperature was 26°C (± 0.5°C), humidity 50% RH (±5%), and circadian light condition was set to 12 hours light/dark cycles (instant switch, no ramping). The maternal generation (F1) was generated in three vial replicates from eggs laid by two sets of F0 flies taken from stock vials. F0 stock vials that were maintained at 21 °C (± 1°C), on the same medium, but with no circadian light control. Stock populations varied between 20 and 60 flies per vial with no larval density control. Nine F0 females and 5 F0 males were selected from stock population, housed together and allowed to lay eggs for 3 days, after which they were discarded. The vials were checked periodically to ensure the development was healthy and progressing at an expected pace. While we set up 3 replicate vials initially, only two were used - the extra vial served as back-up for the cases when a vial needed to be discarded. Any vial showing signs of distress, such as dry food, biofilm, too few larvae or pupae or crowded vials were discarded and the rearing protocol was restarted from the beginning, unless a backup vial was available. This was assessed always by the same experimenter. The F1 progeny were collected 12 days after the last day of egglay. The experimental F2 generation was set up from each vial replicate the same way as the F1 generation. However, after discarding the F1 adult flies, and after 9 days of development, the vials were checked for eclosion daily. As the first few flies eclosed they were discarded. The next day, the peak eclosion would follow, and the newly eclosed flies were collected. Around 40 males and 40 females were then separated into two vials with fresh medium and prepared for the behaviour assay that would follow 3 days later, when the adult flies were 5 days old. All the preparatory flywork, including collection, sorting and counting flies across the generations was performed swiftly and using light CO_2_ anaesthesia (5L/min flow through standard CO_2_ pad) at laboratory room temperature, 21°C (± 0.5°C). All flies were handled by a single experimenter (by R.M. for the single day experiments, and by G.V.B. for multi-day experiments).

### Operant conditioning

We use two operant conditioning paradigms: the associative visual place learning (Figure 1C), and the associative colour learning (Figure 1 - figure supplement 1A). In both paradigms, the unrestrained fly can explore the Y-maze and make sequential binary decisions at the junction of the three arms. Conditioning is initiated after initial measurement of unconditioned behaviour. The decisions during conditioning can result in the absence, introduction or removal of foot shock.

For each genotype, we used two biological replicates, progeny of two different sets of males and females, that were reared independently in different vials. Each replicate consisted of 32 flies, equally split between males and females and between two treatment groups, conditioned and control. Environment during conditioning was kept the same as for rearing. Flies were very lightly anaesthetised (10 L/min CO_2_ flow on 30 x 40 cm custom-made CO_2_ pad) and loaded into the Y-maze arenas using an aspirator. In the conditioned group, flies underwent operant conditioning in the presence of shock, while in the control group the flies were allowed to spontaneously explore the maze experiencing the same visual and environmental stimuli, but without experiencing any shock. All flies were tested during the same “non-siesta” circadian timeframe (20% ZT 0-1, 30% ZT 1-2, 30% ZT 2-3, 15% ZT 3-4) with exception of around 4% of flies extended beyond ZT 3 (3% ZT 4-5, 1% ZT 5-6, 1% ZT 6-7) due to technical issues; Table 7). We found no difference in task performance given the zeitgeber timeframes (ANOVA (Task performance Experiment time (ZT) * Treatment) p = 0.531).

During this time, flies tend to make on average 76 decisions per 20 minutes, with more decisions made by males (N_male_ = 82, N_female_ = 62), and fewer decisions made in the presence of conditioning (N_control_ = 89, N_conditioned_ = 63). In the place learning assay, one of the three arms of the Y-maze is coloured differently from the other two, and an electric shock is associated with that colour. The colours chosen are blue and green, as flies have been shown to be able to distinguish well between the two in previous associative learning studies Ernesto ***Salcedo et al. (1999)***; ***Schnaitmann et al. (2013)***. At the junction of the Y-maze, the fly is presented with a binary choice between two arms. When conditioning to avoid the colour green, the fly gets shocked when it enters a green arm, while the shock is stopped when it leaves the green arm and enters a blue arm. No conditioning occurs when it enters a blue arm from another blue arm. Hence, we are implementing a combination of positive punishment (adding aversive stimulus following incorrect behaviour), and negative reinforcement (removing aversive stimulus following correct behaviour). The shocked arm was associated with a colour assigned to different arms in a clock-wise way, column-wise across the 64 mazes (Figure 1E), and kept fixed for the entire place learning assay. In the colour learning assays the shock arm was associated with a colour, but the colour of the arms changed with every decision, making the shock and the colour uncoupled from a position in space. At the start of any assay, the entire arena is illuminated with red colour for 2 seconds while the floor becomes electrified. This moves the flies from a state of rest to a state of arousal. The video recording is initiated and for the next 3 minutes, the fly is allowed to explore the maze and get accustomed to it without delivering any shock. This time is also used to assess prior individual biases and initial fly behaviours. The operant conditioning assay is then initialized and recorded for 20 minutes.

### Experimental validation of the Y-maze conditioning assay

In the place learning paradigm, wild type conditioned flies spent an average of only 10.5% of the total assay time in the shocked arm, while the control group spent significantly more time in this arm in the absence of shock (T_shocked_ = 32.5 %, Student’s T-test p = 2.52e^-10^). The learning-deficient (*dnc^1^*) showed poor relative task performance at avoiding the shocked arm (conditioned T_shocked_ = 27.8 %, control T_shocked_ = 30.8 %; p = 0.20). Similarly, relative task performance of visually impaired (*norpA^EE5^*) was low (conditioned T_shocked_ = 25.4 %, control T_shocked_ = 25.0 %; p = 0.90; Figure 1F), suggesting that in flies visual context such as colour is informative for place learning ***Whishaw (1998)***. Alternatively, the *norpA^EE5^* mutation may also induce learning impairment beyond perceptual deficiency ***Melnattur et al. (2014)***.

While average genotype performance was consistent across learning paradigms, across different colour stimuli, and, across consecutive measurements of same populations over multiple days, we found that individual learning task performance was variable and evolved over time (Figure 2; Figure 1 - figure supplement 1, Figure 1 - figure supplement 2; Figure 2 - figure supplement 1).

### Multi-day learning task

In the multi-day learning task, genotypes DGRP-362 and *dnc^1^* were reared based on the standard rearing explained above but in 12 replicate vials per genotype (3 replicate vials per experiment type, except for *dnc^1^* for which we had two instead of three replicates for the last two experiments, because two vials were discarded due to poor development). 256 individuals from the two geno-types were collected (total N = 512 flies) to be tested in four types of experiments. Before the test, the collected flies were housed in fly vials in groups of around 40 individuals of mixed sexes. On the day of each first test, 32 females and 32 males were selected for the test. In each of the four experiments, a set of 64 individual flies were tested once per day with the exception of the second experiment of DGRP-362 that contained 59 flies (32 males and 27 females) due to experimenter error (fly identities were accidentally and intractably swapped). In total N = 507 flies were tested successfully. Flies were tested on two consecutive days followed by one day without a test, then another set of two test days. All tests were done under conditioned treatment. Before each conditioning session, flies were allowed to explore the arena for 5 minutes in the absence of shock. A 20-minute conditioning session ensued, followed by a 5-minute session in the absence of shock, to assess change in bias due to learning and short-term memory. This regime was repeated once per day, at the same time of the day (Zeitgeber 0-2). In the first experiment (PPCC_gbgb), flies were tested in a place learning paradigm the first two days (”PP”), then in the colour learning paradigm the last two days (”CC”). The colours associated with the shock across four days alternated across tests between green and blue (”gbgb”). In the second experiment (PPCC_bgbg), they alternated between blue and green. In the third (CCPP_gbgb) and fourth experiment (CCPP_bgbg), on the first two days place learning was tested, and on the last two days, we tested the flies in a colour learning paradigm. The colours associated with the shock alternated the same way for the third and fourth experiment, as in the first and second, respectively. Between tests, identity of each fly was tracked by housing them individually in a custom 3D-printed 64-well single housing plate, placed on food and isolated with a dome to avoid the food from drying up. The flies were transferred to the single housing upon completion of the each test. Each fly was always tested in the same Y-maze on the platform over the days. The position in the well plate corresponded to the position on the platform to facilitate quick set up of the experiment. Males were placed on the top 4 rows of the platform and females on the 4 bottom ones. All flies were handled by a single experimenter (by G.V.B.), except for the third experiment (where the assay was set up by A.M.J.).

### Single fly housing

The single housing plate used in the multi-day learning task consists of 64 cube-shaped wells, organized in 8 rows and columns. The floor of the plate is a 0.2 mm thick printed mesh. The mesh allows the flies to reach food and eat when the plate is placed on a tray filled with fly medium, while keeping them as dry and clean as possible throughout the experiment. Care should be taken to only gently press the well-plate on the surface of the medium and prevent food from entering the wells. The mesh also helps with anaesthetising the flies by simply placing the plate onto the CO2 pad. The top of the 64-well plate is covered with a 3D-printed lid, shaped as an inverse imprint of the 64-well plate. This way, when it covers the well plate, the grid of the lid protrudes by 0.2 mm into each well, preventing the flies from wiggling their way between the lid and the plate and crossing into another well. A thin, fine filter mesh, usually used for Drosophila egg collection, was printed into the top printed layers of the lid. To manufacture this, the top four 0.06 mm layers of the lid are printed first. Then, the printer is paused, the filter mesh is laid over the printed part and tightly affixed to the printer’s build plate with an adhesive tape. The printing is resumed so that the next layers of print material “bakes” the mesh into the print. After printing, the mesh remaining on the outer part of the print is cut with a precision knife. The .stl models can be found in the github repository. The plate and lid were printed using red PLA filament (to avoid colour imprinting or colour bias) on an Ultimaker S5 3D-printer. Black filament can be used as well, but the low contrast with the fly makes picking and placing flies from the well more challenging and prone to errors. The 3D models were sliced with Ultimaker Cura, using default “Fine resolution” print setting (0.06 mm layer height). To create the bottom printed mesh, we use Ultimaker Cura slicer to slice a 0.2 mm thick block with width and length matching the size of the dimensions of the 64-well plate. We use the 60% “grid” infill and omit the bottom and the top layer print. Once printed, the mesh is glued to the bottom of the 64-well plate using cyanoacrylate adhesive. This has to be prepared at least 24 hours before the experiment, in a well-ventilated room, to allow the volatile compounds of the adhesive to fully air out. Occasional slight warping of the lid would make it possible for flies to cross to other wells, thus losing track of their identity. Thus, testing the lids before an experiment is advised.

### Basic statistics

All basic statistical tests were performed within the R environment (R version 4.0.2.). Pearson correlations and T-tests were performed using base cor.test() and t.test() functions. T-tests were two-sided. Resampling test was performed using the function lmperm() from the package permuco ***Frossard and Renaud (2021)***. Exact general independence test was performed using the function independence_test() from the package coin ***Strasser and Weber (1999)***; ***Hothorn et al. (2008)***. Generalized linear mixed models were fitted using lmer() function from the lmerTest package and the analysis of variance for fitted models was performed using the anova() function. Least square means and the pairwise difference between them were calculated using emmeans (pbkrtest.limit = 100000000) and pairs() function from lsmeans R package. Where multiple tests were compared p-values were corrected using Bonferroni correction for multiple test using base p.adjust(method = “bonferroni”) function. Density distributions are estimated as kernel densities using density() function and default parameters.

### Definitions of behaviours and other behaviour-based measures

*Task performance* (T_shocked_) of a fly is the fraction of time a fly has spent in the conditioned arm during the length of the experiment (20 minutes). It is measured the same way regardless whether the shock is applied or not (for example in the control experiment). *Handedness* of a fly is the absolute value of the total number of right turns subtracted from the total number of left turns, divided by the total number of turns taken by the fly at any given time frame. *Developmental noise* is the absolute handedness of a fly measured during the three minutes prior to the conditioning. *Velocity* in mm/s for a given time frame, is calculated by averaging the displacement of a fly per second over the said time frame. *Relative velocity* is the difference in average velocity of a fly between the shock-associated arm of the Y-maze and the neutral arm of the Y-maze at a given time frame. *Shock response* is the absolute value of the difference between the average velocity of a fly measured during the three minutes prior to the start of the conditioning part of the experiment and the average velocity of the fly during the first quarter (5 min) of the conditioning experiment (when shock becomes implemented in the conditioned group). *colour bias* of a fly is defined as the total fraction of the time the fly spends on the colour during a set time frame. *Percent correct decisions* is the proportion of made decisions to enter the non-shocked arm out of all made decisions to enter any arm. *Learning score* is computed by splitting the experiments into 10 bins of 2 minutes and computing the negative of the average of two Spearman’s rank correlations: 1) the rank correlation of the interval index *𝑖* with the relative time spent in the shock arm during interval *𝑖* and, 2) the rank correlation of the interval index *𝑖* with the relative number of visits to the shock arm during interval *𝑖*.

### Genotype-specific relative task performance

Before calculating mean task performance for a genotype, we first filter out individual flies based on their lack of activity. Flies are removed from the analysis if they remain at the shocked arm for 3/4 of the experiment or if they never enter the shocked arm/colour (in either conditioned and control group of flies). Based on observations in trial and final experiments, this outlier behaviour was due to the fly getting accidentally injured by mis-manipulation during assay setup (usually by slightly misplacing them outside the maze boundary and pinching them with the glass slide). The filtering accounted for 3.78 % of all tested wild-type flies. A linear mixed modelling is then used to estimate the effect of genotype on the response to shock and to regress out nuisance factors that may affect the measures of learning. Those included sex, replicate, sex-specific locomotor activity during and in the absence of shock, developmental noise, position of the Y-maze, position of the shocked arm, position of the arm in which the fly initializes the task. The model with the T_shocked_ as the response variable, genotype interacting with the treatment (conditioned or control), sex, developmental noise (absolute bias locomotor handedness), shock response and the starting arm (shock or neutral left, or neutral right) as the fixed effects, and the replicate number, and the position of the fly on the 64-maze arena as the random effects, is fitted to the data to capture individual components of variance of the data (Table 9,10). The least squared means of the genotype x treatment effect was extracted. Relative task performance of a genotype was measured as the Tukey’s Honestly Significant Difference in least square mean of T_shocked_ for a genotype in the conditioned group and least square mean of T_shocked_ for a genotype in the control group.

### Individuality

Extent of expressed individuality in a behaviour within a group of flies is measured using the differential entropy ***Cover and Thomas (2006)*** of the probability distribution function of the behaviour for that particular group, estimated over the theoretical range of the behaviour (Table 12). For Gaussian distributions, entropy scales with the logarithm of the standard deviation and is thus consistent with classical measures of variability in isogenic populations, often called individuality (Figure 2 - figure supplement 8). The difference in expressed individuality between two groups of flies is calculated as Hellinger distance ***Hellinger (1909)*** between the two distribution functions. Specifically, we calculate Hellinger distance between two continuous probability density functions of the distributions, over the theoretical range of the concerned behaviour.

### Residual individuality and distribution shape divergence

To assess the extent of residual individuality (H_resid_), the distributions of expressed individual behaviours within genotypes, groups, or replicates are first independently divided into a histogram of 10 bins within the individual range specific to each distribution. The probabilities of finding a measurement in each of the 10 bins are then calculated to obtain the binned probability distribution, based on which the Shannon entropy ***Shannon (1948)***; ***Hausser and Strimmer (2008)*** is calculated. The H_resid_ correlates negatively with the range of T_shocked_ (Figure 2 - figure supplement 8). This is a result of scaling the distributions within their own ranges, and is consistent with our definition of H_resid_ being free from the effects of environmental and developmental stochasticity. That is, assuming we are able to capture the full range of the expressed behaviour (evidenced by consistency of the distributions between replicates), a group of flies having a narrow range of T_shocked_, will be scaled to an approximately uniform distribution within the range of the particular distribution (Figure 2 - figure supplement 6), and hence, have a higher entropy. To capture the difference in expressed residual individuality between two groups, the same independent binning approach is taken for each group-specific probability distribution separately, and the Kullback-Leibler (KL) Divergence ***Kullback and Leibler (1951)***; ***Drost (2018)*** of the two distributions is calculated. Since each distribution is binned independently within their own range, every distribution becomes a list of 10 probability values (summing up to 1), with each value having an equal weight for the calculation of the entropy as well as the divergence. This procedure scales each probability distribution to a common range (Figure 2 - figure supplement 6), and ensures that the entropy and the divergence measures are only capturing the shapes without being confounded by the central tendencies or the ranges of the distributions. Effectively, we capture residual individuality that is expressed independently of the variability caused by developmental stochasticity or environmental plasticity. The KL divergence is a non-metric, asymmetric measure of how different one distribution is from a reference distribution. Therefore, when we measure the change in the residual individuality as a result of conditioning (*Learning divergence*, Δ *H_resid_ Learning*), we measure the KL divergence of the conditioned (shocked) population with respect to the control (non-shocked) population. In other cases where no reference distribution is obvious, KL is calculated in both directions (such as in the comparison between genotypes within a treatment, *genetic divergence*, Δ *H_resid_ Genetic*), or the compared distribution pairs are randomized, and the pairs are kept consistent between tested environments (such as in the comparison between replicates within a genotype and treatment, *experimental divergence*, Δ *H_resid_ Experimental*).

To provide an alternative measure of differences in the shapes of distributions that are symmetric or less sensitive to outlier values we also calculated Hellinger distance ***Hellinger (1909)*** and Jensen-Shannon divergence J. ***Lin (1991)*** between the distributions of T_shocked_ in control and conditioned treatment, that were centered around the median. We also applied the KL divergence on non-binned, continuous distributions of T_shocked_ (Figure 2 - figure supplement 8). We found that these measures all correlate strongly and significantly with each other and with either the range, variance and entropy of the T_shocked_ distributions and/or the relative task performance. While they do uncover various differences between distributions, these measures are not free from variance and central tendency effects. Hence, we refrain from using them for measuring the changes in *Residual individuality*.

### Modeling behaviour

To model individual fly behaviour, we extract for each fly a sequence of arm identities *𝑥_𝑖_* ∈ {1, 2, 3}, arm colour *𝑐_𝑖_* ∈ {green, blue}, presence of shock *𝑠_𝑖_* ∈ {shocked, not shocked}, arm escape-duration Δ*𝑡_𝑖_* ∈ ℝ^+^ and turn events *𝑦_𝑖_* ∈ {right, lef t}. For example, the sequence *𝑆* = ((*𝑥*_1_ = 3*, 𝑐*_1_ = green*, 𝑠*_1_ = shocked, Δ*𝑡*_1_ = 1.0 s*, 𝑦_𝑖_* = right), (*𝑥*_2_ = 1*, 𝑐*_2_ = blue*, 𝑠*_2_ = not shocked, Δ*𝑡*_2_ = 3.5 s*, 𝑦*_2_ = lef t), …) means that a fly started in arm 3, perceived colour green and got shocked, escaped arm 3 after one second by taking a right turn at the center of the maze, arrived in arm 1 with colour blue and without shock, stayed there for 3.5 seconds and left it with a left turn, etc. We model the probability of a given sequence *𝑆* of length *𝑇* as 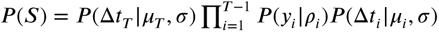, where *𝑃* (*𝑦* |*𝜌* ) is a Bernoulli distribution with probability of turning right given by *𝜌_𝑖_* and *𝑃* (Δ*𝑡_𝑖_ 𝜇_𝑖_, 𝜎*) is a log-normal distribution with mean *𝜇_𝑖_* and standard deviation *𝜎*. The probability of turning right 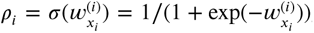, depends on the weights 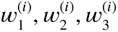 (one for each state). These weights change according to the following dynamics: 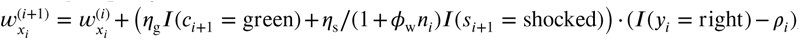, where *𝜂*_g_ is the green colour learning rate, *𝐼* denotes the indicator function with *𝐼* (true) = 1 and *𝐼* (false) = 0, *𝜂*_s_ is the shock learning rate, *𝑛_𝑖_* is a shock counter (number of times shock was experienced so far), and *𝜙*_w_ is a parameter that scales the dependence on the shock counter. This learning rule increases the probability of turning right in state *𝑥_𝑖_*, when turning right led to the green arm without shock and *𝜂*_g_ *>* 0 (colour preference learning), and it leads to a decrease in the probability of turning right, when the negative impact of arriving in a shocked arm outweighs the preference for arriving in a green arm (*𝜂*_g_ + *𝜂*_s_∕(1 + *𝜙𝑛_𝑖_*) *<* 0), and vice versa for turning left. The means of the log-normal distribution are given by *𝜇_𝑖_* = *𝜇*^(^*^𝑖^*^)^ + Δ_g_*𝐼* (*𝑐_𝑖_* = green) + Δ_s_*𝐼* (*𝑠_𝑖_* = shocked), where the parameters Δ_g_ and Δ_s_ control the change in escape duration due to being in a green arm or being shocked, relative to the mean *𝜇*^(^*^𝑖^*^)^. This mean changes according to the dynamics *𝜇*^(^*^𝑖^*^+1)^ = *𝜇*^(^*^𝑖^*^)^ + *𝜂_𝜇_* ∕(1 + *𝜙_𝜇_𝑛*_s_)*𝐼* (*𝑠_𝑖_* = shocked) + *𝜂*_t_ Δ*𝑡_𝑖_*. With this update rule, the mean of the log-normal distribution over escape durations can change simply with the passage of time, when *𝜂*_t_ ≠ 0, but also due to being shocked, when *𝜂_𝜇_* ≠ 0. To model a population of flies, e.g., 64 flies of the same genotype, we place normal priors over the model parameters (Table 13) and estimate the means and standard deviations of these priors with approximate expectation-maximization. We considered four different models: no individuality: all flies with the same genotype have exactly the same parameters, i.e., the standard deviation of all priors is zero (*𝑘* = 11 parameters per genotype; (Table 13).

**Table 13.** Model parameters. The first five parameters are “static preference” parameters and the remaining parameters are “learning rate” parameters.

| symbol | description |
| --- | --- |
| $w^{(1)} \in \mathbb{R}$ | initial probability of turning right (handedness) |
| $\mu^{(1)} \in \mathbb{R}$ | initial mean of the log-normal distribution over escape durations |
| $\Delta_s \in \mathbb{R}$ | shock-induced change of the mean of the log-normal distribution |
| $\Delta_g \in \mathbb{R}$ | change of the mean of the log-normal distribution, because of being in a green arm |
| $\sigma \in \mathbb{R}^+$ | standard deviation of the log-normal distribution over escape durations |
| $\eta_s \in \mathbb{R}$ | learning rate controlling the shock-induced change of turning right |
| $\eta_g \in \mathbb{R}$ | learning rate controlling the green-color-induced change of turning right |
| $\eta_\mu \in \mathbb{R}$ | learning rate controlling the shock-induced change of the mean of the log-normal distribution over escape durations |
| $\eta_t \in \mathbb{R}$ | learning rate controlling the change of the mean of the log-normal distribution, because of the passage of time |
| $\phi_w \in [10^{-3}, 10^3]$ | factor scaling the influence of the shock counter on the turn dynamics |
| $\phi_\mu \in [10^{-3}, 10^3]$ | factor scaling the influence of the shock counter on the escape duration dynamics |

individuality: normal priors with fitted mean and standard deviation over all parameters, except the learning rates *𝜂*_s_*, 𝜂*_g_*, 𝜂_𝜇_* and the factors *𝜙*_w_*, 𝜙_𝜇_*, which are shared across individual flies of the same genotype, as in the no individuality model (*𝑘* = 11 + 6 = 17 parameters per genotype). Sharing the learning rates led to a lower BIC than a model with individuality in all parameters (not shown).

no learning: same as individuality, except that *𝜂*_s_ = *𝜂*_g_ = *𝜂_𝜇_* = *𝜂_𝑡_* = 0 and *𝜙*_w_ = *𝜙_𝜇_* = 1 (*𝑘*= 11+5−6 = 10 parameters per genotype).

no genetics: all flies are fitted with the individuality model (*𝑘* = 17 parameters), shared parameters due to genotype is not modeled.

Bayesian information criterion BIC and marginal log probabilities were estimated with 10^4^ samples. The difference in momentary behaviour between two simulated flies was computed with the Hellinger distance ***Hellinger (1909)*** between the policies, averaged over all possible states.

### Simulating behaviour

We simulated tracks for 64 flies per genotype split equally between treatments based on the genotype-specific or individually fitted parameters from the different models.

### Predicting genotype from behaviour

To decode the genotype from behaviour, we trained a recurrent neural network with one hidden layer of 32 LSTM cells with relu-activation on the behavioural sequences encoded as (onehot(*𝑥_𝑖_*)*, 𝑠_𝑖_,* Δ*𝑡_𝑖_*). The feed-forward baseline model contained 2 hidden layers of 32 relu neurons with the input being the number of decisions encoded as 20 dimensional vectors with Gaussian tuning, i.e., for number of decisions *𝑛*, input neuron *𝑖*’s activity is given by *𝑎_𝑖_* = exp(−(*𝑛* − *𝑐_𝑖_*)^2^∕(2 ⋅ 10^2^)) with *𝑐_𝑖_* = (*𝑖* − 1) ⋅ 20 + 2.

## Acknowledgments

We would like to thank Samuel Bourgeat for helpful discussions and help with fly care, Wulfram Gerstner for the compute time on the cluster, the PCB Lab at the EPFL’s School of Engineering, for providing equipment and material for the fabrication of printed circuit boards used in the project, and the members of the “Project Flybot” WhatsApp chat group (Tomislav Štampar, Stjepan Bukal, Marko Jakšić and Samuel Bourgeat) for the moral and technical support during design of the plat-form.

## Author contributions

AMJ and RM conceptualized and designed the study. RM, AMJ and IT conceptualized and designed the behaviour platform. RM built the platform’s software, RM and AMJ built the experimental hardware. RM, GVB and AMJ performed the experiments. RM, JB, AMJ and AM conceptualized the data analysis. RM, JB, GVB and AMJ analyzed the data and produced the results. AMJ, RM and JB wrote the manuscript. All authors read, edited and intellectually significantly contributed to the manuscript.

## Supplementary Figures and Tables

**Figure 1—figure supplement 1.**
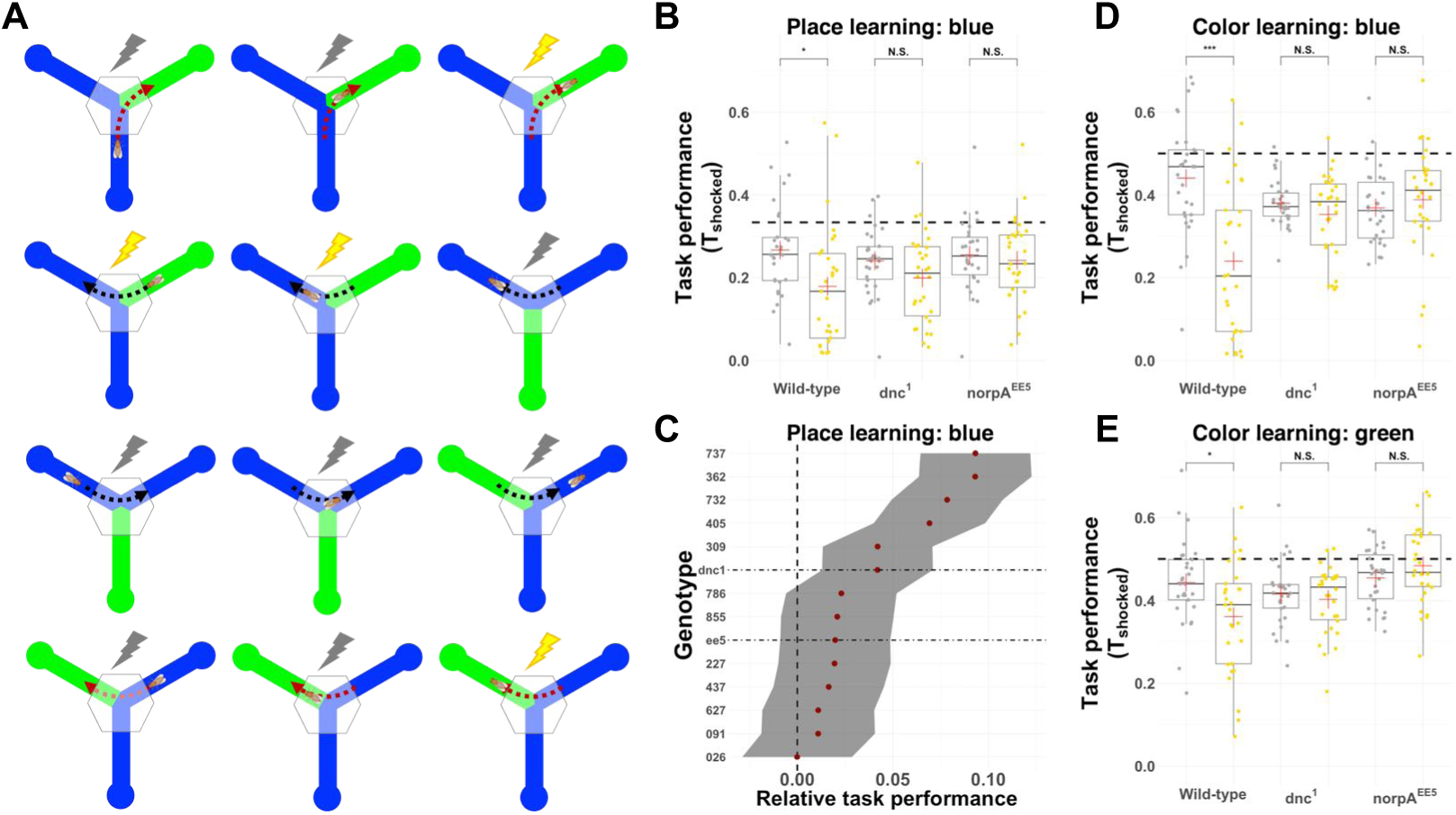
A) Schematic for the implementation of colour learning paradigm. As the fly enters the arm with shocked colour (green) and traverses the 2 mm boundary, it starts to experience the shock. At this point, the decision is recorded and with each decision the colours of the two unoccupied arms behind the fly randomly change colours. This way, the shock is associated with a colour and uncoupled from place. As the fly chooses the neutral colour (blue) the shock is removed (or not applied). B) Task performance in blue place learning for one wild-type DGRP line (DGRP-362), memory deficient genotype (dnc^1^) and blind genotype (norpA^EE5^). Each point is an individual fly. Control treatment is shown in grey and conditioned treatment in yellow. Dashed line marks the T_shocked_ = 1/3. C) Relative task performance in blue place learning for 12 wild-type DGRP genotypes, memory deficient genotype (dnc^1^) and blind genotype (norpA^EE5^). Grey shading indicates standard error. Vertical line points to no change in performance (0) and horizontal lines inidcate least square means for memory- and vision-deficient genotypes. In all panels for each genotype-treatment combination N = 32 flies were tested, but fewer are shown here (due to filtering for low activity; see Methods). For all boxplots: whiskers of boxplots show upper and lower quantiles, the bar indicates median and red plus indicates mean across individuals (points), yellow indicates the conditioned and grey the control flies, asterisk indicates significance threshold for the T-test (* *<* 0.05,** *<* 0.01, *** *<* 0.001), and horizontal lines indicate task performance expected by chance (1/3 or 1/2). D) Task performance in blue colour learning for one wild-type DGRP line (DGRP-362), memory deficient genotype (dnc^1^) and blind genotype (norpA^EE5^). Each point is an in-dividual fly. Control treatment is shown in grey and conditioned treatment in yellow. Dashed line marks the T_shocked_ = 1/3. E) Task performance in green colour learning for one wild-type DGRP line (DGRP-362), memory deficient genotype (dnc^1^) and blind genotype (norpA^EE5^). Each point is an individual fly. Control treatment is shown in grey and conditioned treatment in yellow. Dashed line marks the T_shocked_ = 1/3.

**Figure 1—figure supplement 2.**
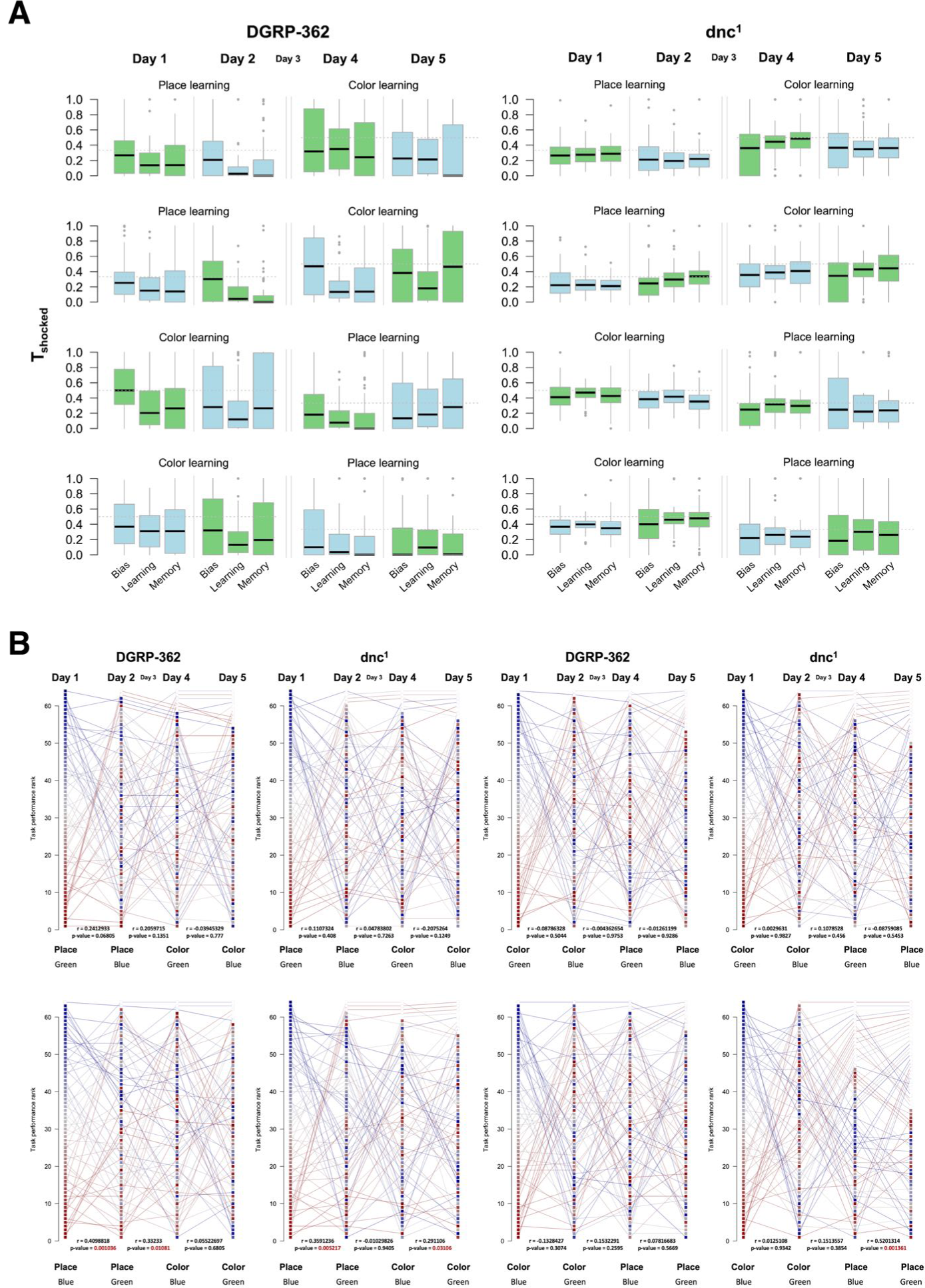
A) Task performance across individuals is shown as boxplots, each for bias period of 5 minutes before conditioning where no shock is applied (”Bias”), 20-minute conditioning period (”Learning”) and 5-minute period where no shock is applied used to assess short-term memory based on the persistence of task performance (”Memory”). Shock was associated with the colour matching the colour of the boxplot (blue or green). B) Individual flies from the same experiment in A) are ranked based on their task performance during conditioning period on the first day from highest (red) to lowest (blue) T_shocked_. Their task performance rank during conditioning period is tracked across consecutive days and paradigms. If the flies died or were lost during the experiment, their past relative ranking is depicted as the line, but the square is not shown for the days after they were lost. Pearson product-moment correlation (r) between ranking of flies on consecutive days is shown. The associated statistical significance of the correlation (p-value) is coloured red if it had passed the FDR corrected threshold of FDR *<* 0.05. For all boxplots, whiskers show upper and lower quantiles, the bar indicates median.

**Figure 2—figure supplement 1.**
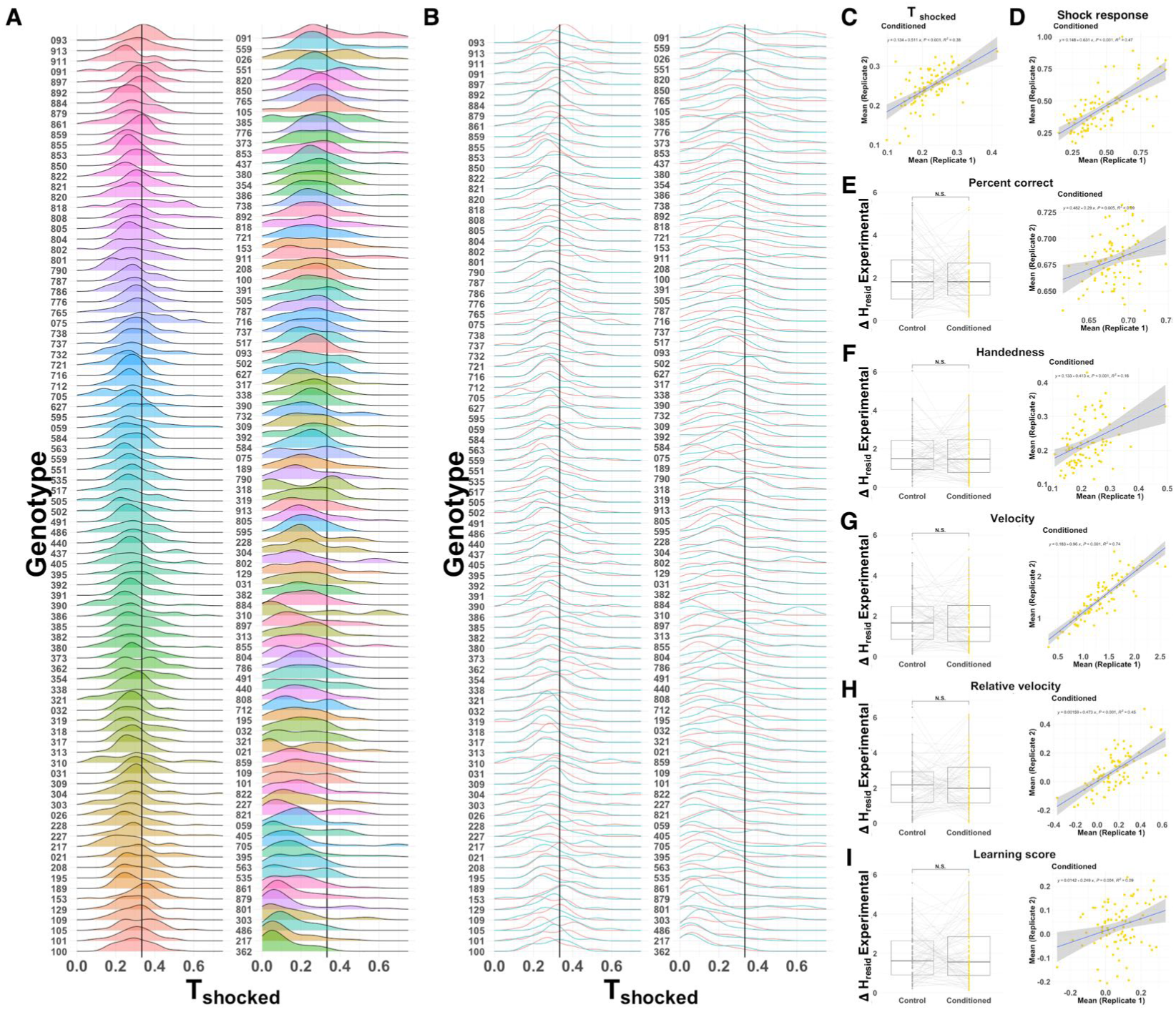
A) Density distributions of task performance for genotypes, or-dered from highest mean T_shocked_ (top) to lowest mean T_shocked_ (bottom) for control (left panel; N = 32 for each genotype) and conditioned (right panel; N = 32 flies for each genotype) treatment. Distribution filled colours represent different genotypes. Solid vertical black lines represent chance level T_shocked_ = 1/3; B) Density distributions of replicates for genotypes shown in A); C-D) Correlation between replicates of mean task performance (C) and shock response (D) ( N = 16 flies per replicate per genotype for 88 genotypes). Inset text show correlation equation, p-value and R^2^; E-I). Δ *H_resid_ Experimental* in behaviour for control and conditioned flies across genotypes (N = 88 genotypes, N.S. indicates no significant difference (T-test) between tested groups; left panel); and correlation of mean behaviour between replicates ( N = 16 flies per replicate per genotype for 88 genotypes; right panel). Inset text show correlation equation, p-value and R^2^; Same is shown in E-I) where E represents correct decisions, F) handedness, G) velocity (mm/s), H) relative velocity, and I) learning score.

**Figure 2—figure supplement 2.**
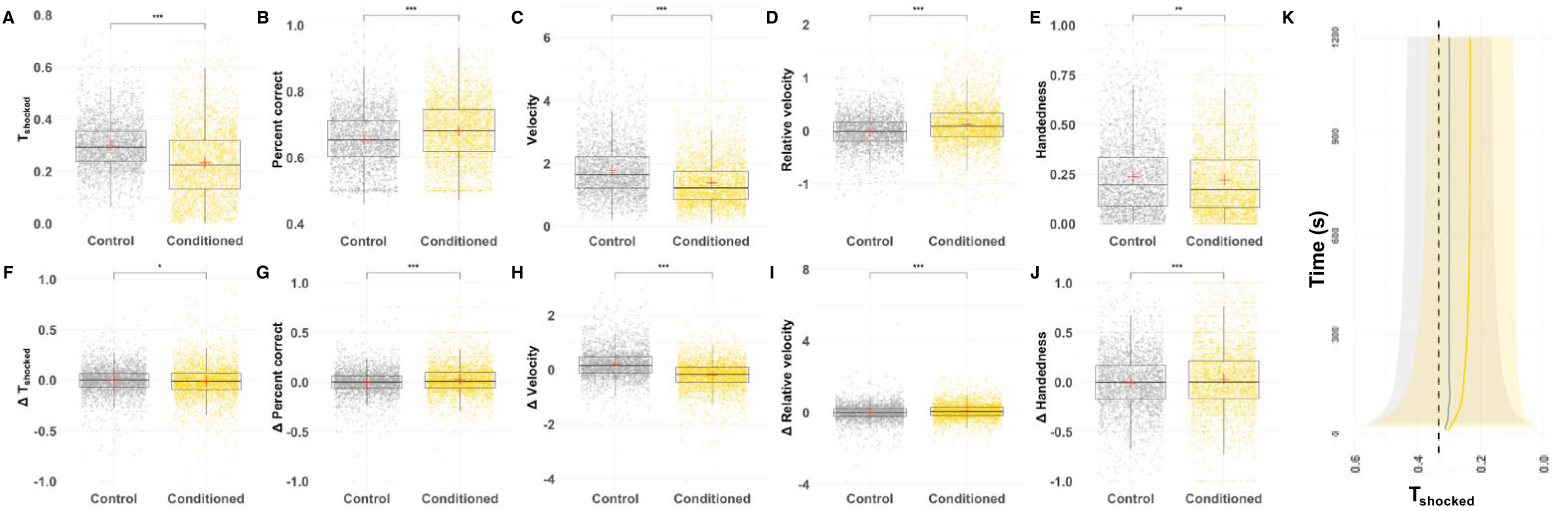
A-E) Individual fly behaviour and change in behaviour in condi-tioned (yellow) and control (grey) treatments is shown for A) task performance, B) Percentage of correct decisions, C) velocity (mm/s), D) relative velocity (mm/s) and E) handedness. F-J) Change in individual behaviour between 4th quarter (15 - 20. min) and 1st quarter (0 - 5 min) of the condition-ing assay is shown for the same behaviours. K) Evolution of task performance over 20-min assay. Solid yellow and grey lines show the average cumulative task performance for all conditioned flies (N = 2611) and control flies (N = 2628), respectively. Shaded areas show 1 standard deviation. For all boxplots, whiskers show upper and lower quantiles, the bar indicates median and red plus indi-cates mean across individuals (points). In all panels yellow indicates the conditioned and grey the control flies. Asterisk indicates significance threshold for the T-test (* < 0.05,** < 0.01, *** < 0.001)

**Figure 2—figure supplement 3.**
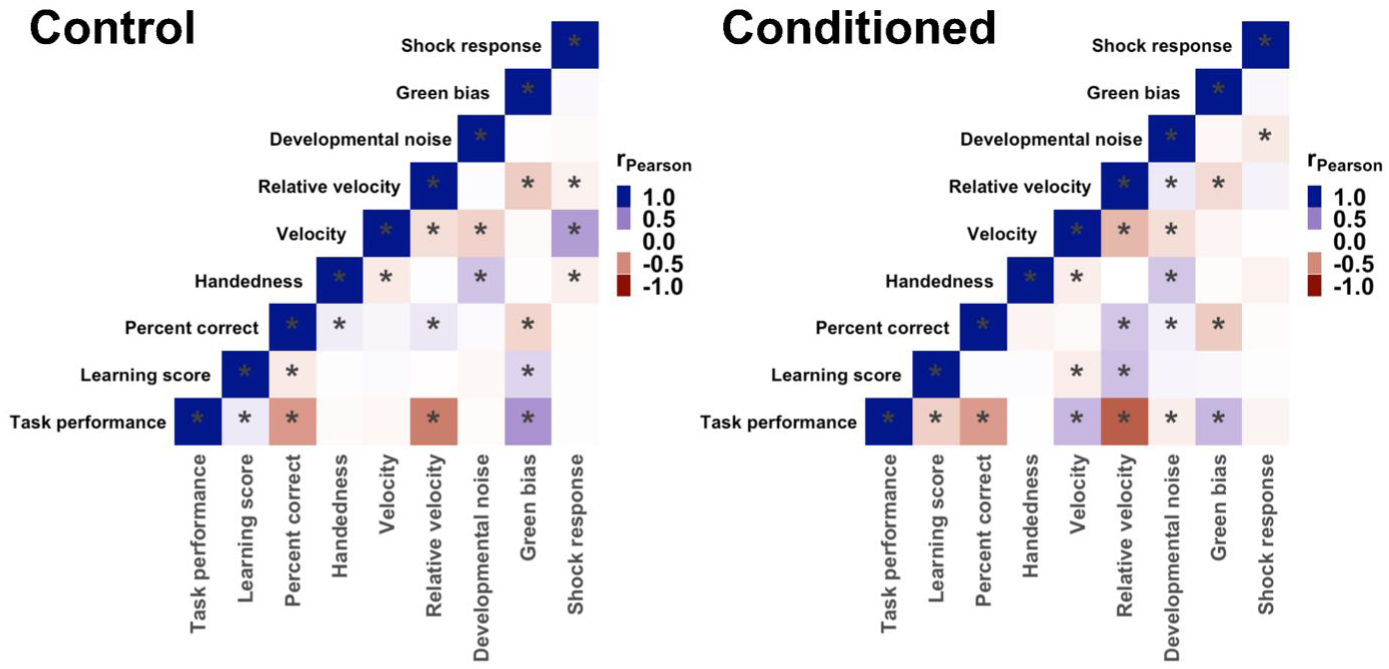
Heatmap shows Pearson’s product moment correlation matrix between individual’s behaviours in the conditioned and control treatment. Pearson product-moment correlation coefficient (r) scale: Blue r = 1, white r= 0, red r = -1. Asterisks indicate p-value < 0.05 after Bonferroni correction for multiple testing. For all boxplots, whiskers show upper and lower quantiles, the bar indicates median. In all panels yellow points indicate the conditioned and grey the control flies. Asterisk indicates significance threshold for the T-test (* < 0.05,** < 0.01, *** < 0.001).

**Figure 2—figure supplement 4.**
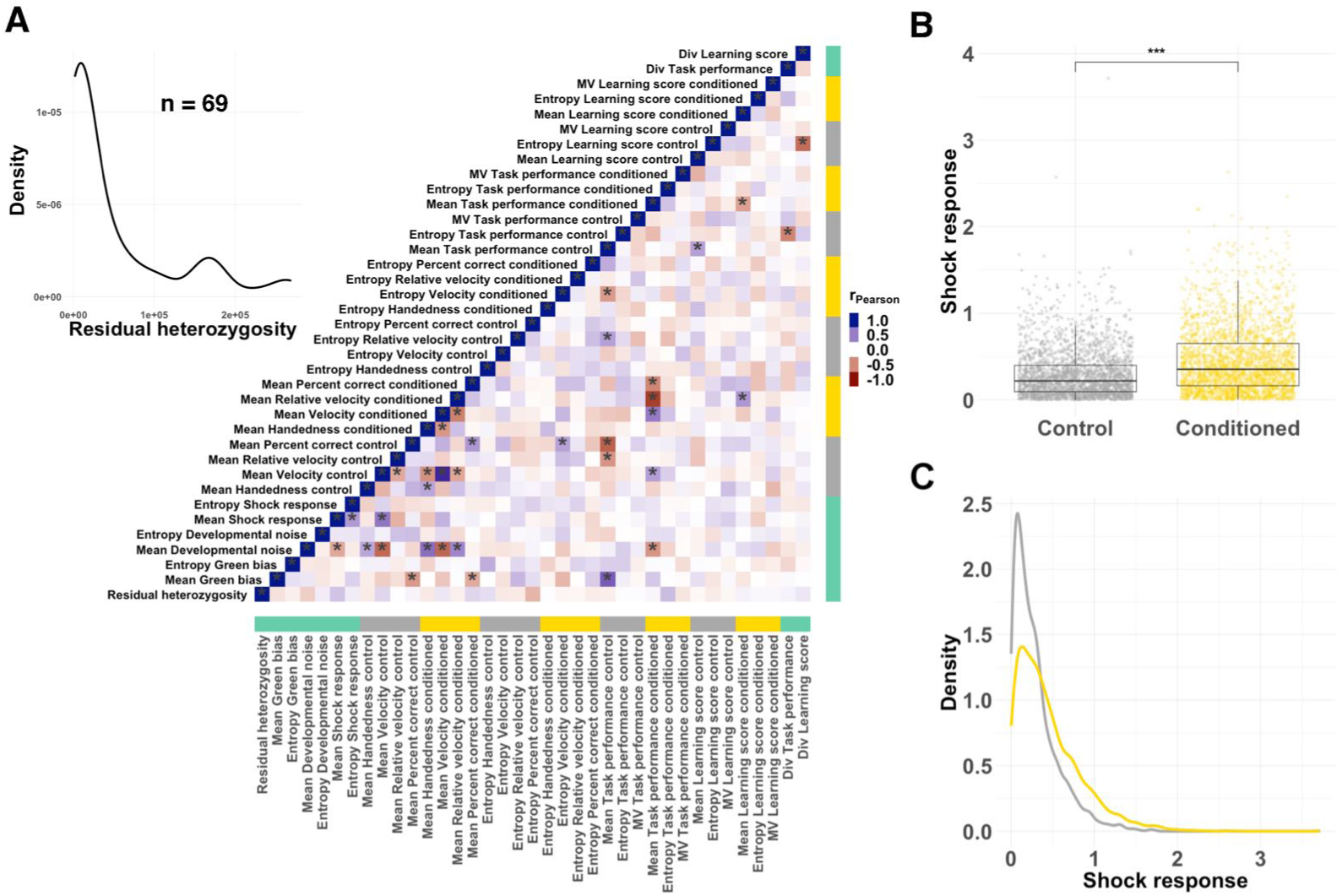
A) Correlation matrix for genotype scores of various measured fly behaviours (N = 88 genotypes). Yellow bands indicate the conditioned groups, grey bands the control groups, green bands represents scores measured over all flies regardless of the treatment. Pearson product-moment correlation coefficient (r) scale: Blue r = 1, white r= 0, red r = -1. Asterisks indicate p-value < 0.05 after Bonferroni correction for multiple testing. Inset shows the distribution of residual heterozygosity (total number of heterozygous loci) for 69 genotypes as reported in ***Mackay et al. (2012)***; B) Shock response for control (grey, N = 2628) and conditioned flies (yellow, N = 2611). For all boxplots whiskers of boxplots show upper and lower quantiles, the bar indicates median across individuals (points). Asterisk indicates significance threshold for the T-test (* < 0.05,** < 0.01, *** < 0.001). C) Kernel density distribution of shock response for control (grey) and conditioned (yellow) flies.

**Figure 2—figure supplement 5.**
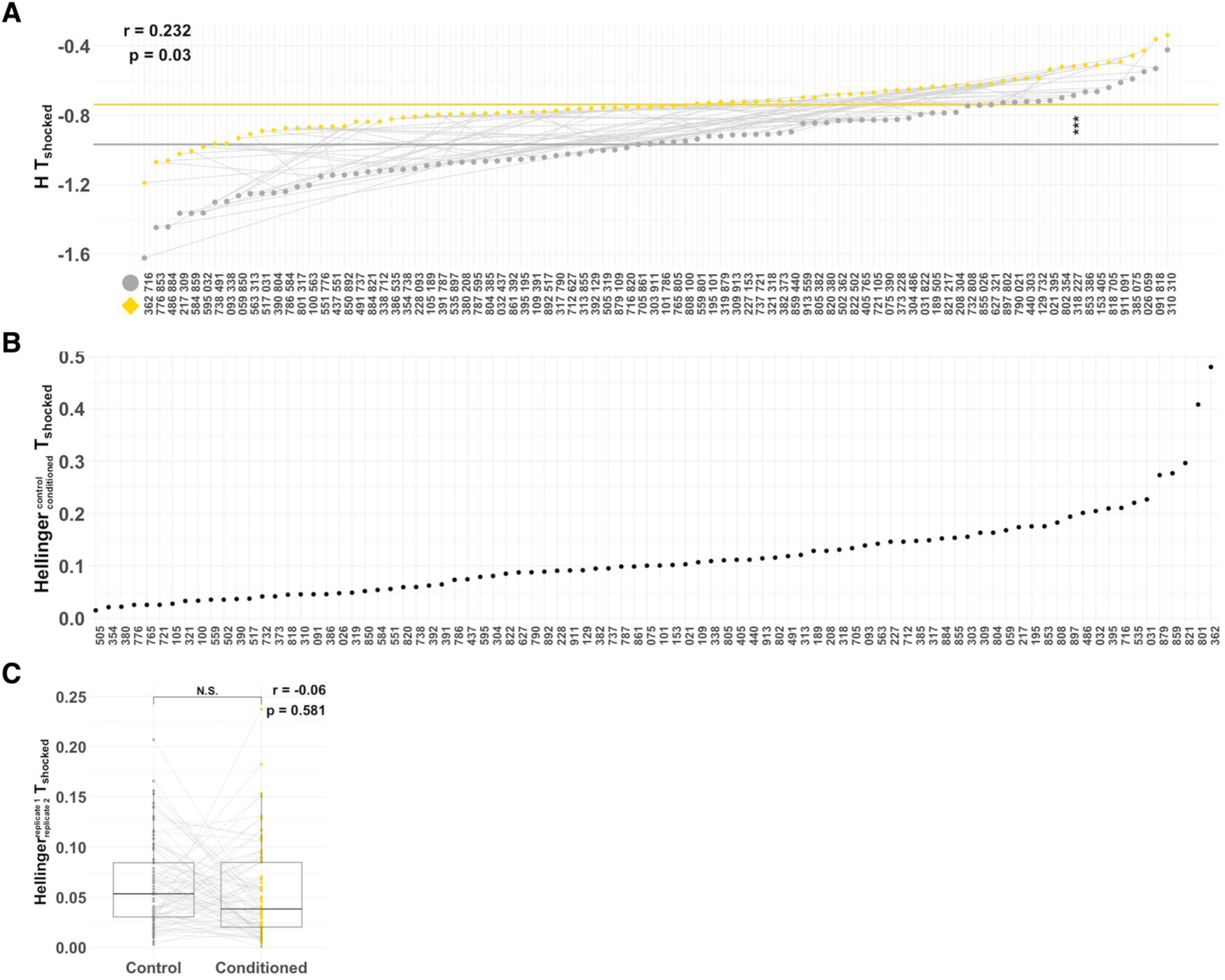
A) Task performance entropy (*H*) across 88 wild-type genotypes measured in the control (gray) and conditioned (yellow) treatment. Genotypes are sorted independently for control and conditioned treatment. Thin gray lines connect the same genotype across two treatments. Horizontal lines indicate mean entropy value across genotypes. Asterisks indicate a significant difference between the means across genotypes. B) Hellinger distance between the task performance distributions in control and conditioned flies is shown for each genotype. Geno-types are ranked based on the Hellinger distance, from lowest to highest. C) Task performance distributions are similar in replicate 1 and replicate 2.

**Figure 2—figure supplement 6.**
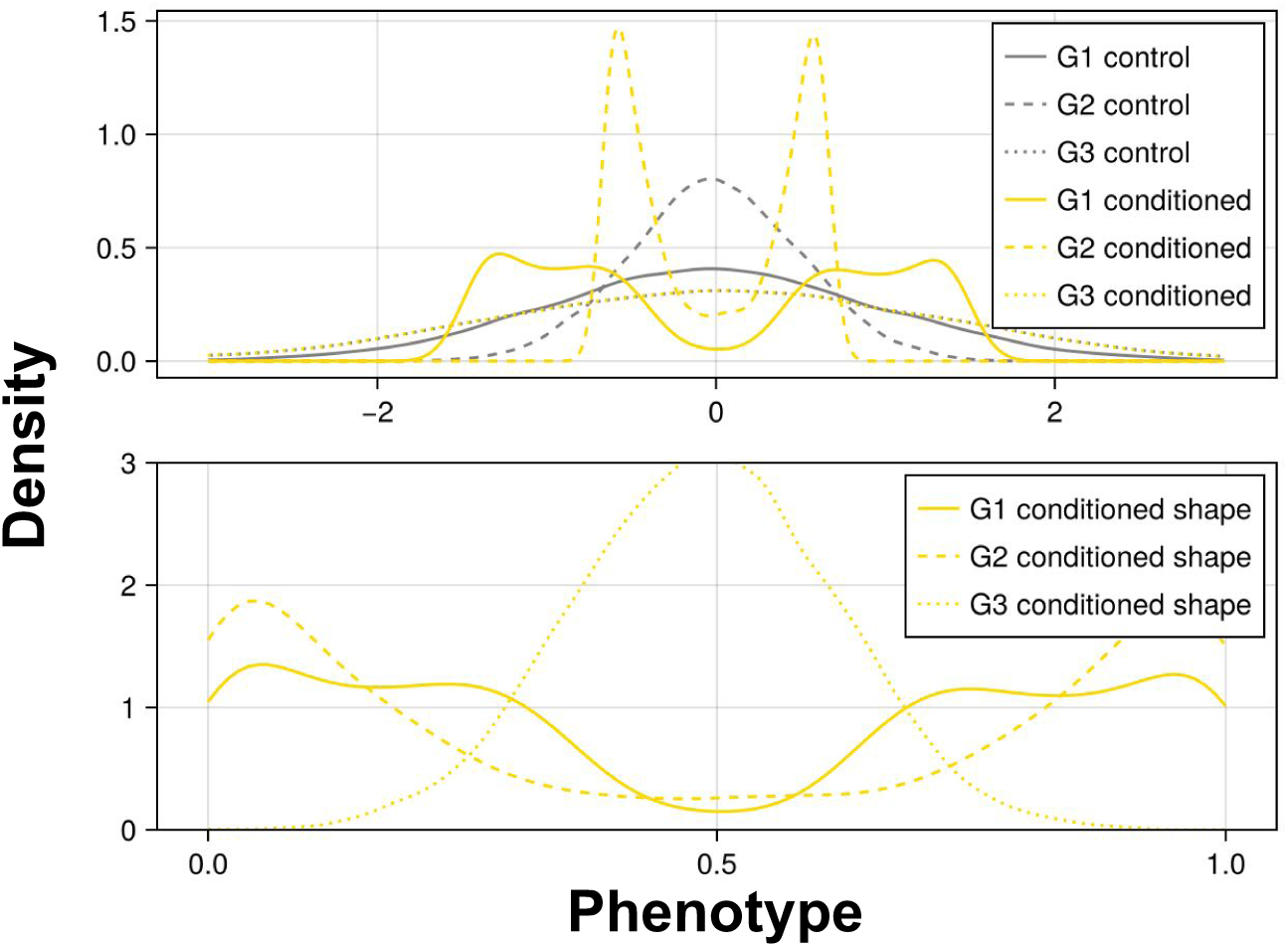
A schematic representation of how scaling distributions to the same range can be used to assess the shape of the distribution independent of the distribution variance. Assume some measurements with genotypes G1, G2 and G3 have a normal distribution under control condition (top figure, gray lines). In the conditioned case, the distributions are bimodal for G1 and G2, whereas it remains normal for G3 (top panel). For each genotype, the variance is the same in both conditions. If we simply compute the Hellinger distances, we find D(G1 conditioned, G2 conditioned) > D(G1 conditioned, G3 conditioned), because the Hellinger distances are dominated by the variance here and G1 and G3 have similar variances. However, when we transform the measurement variables to the same range [0, 1] (bottom panel), we find D(G1 conditioned shape, G2 conditioned shape) < D(G1 conditioned shape, G3 conditioned shape).

**Figure 2—figure supplement 7.**
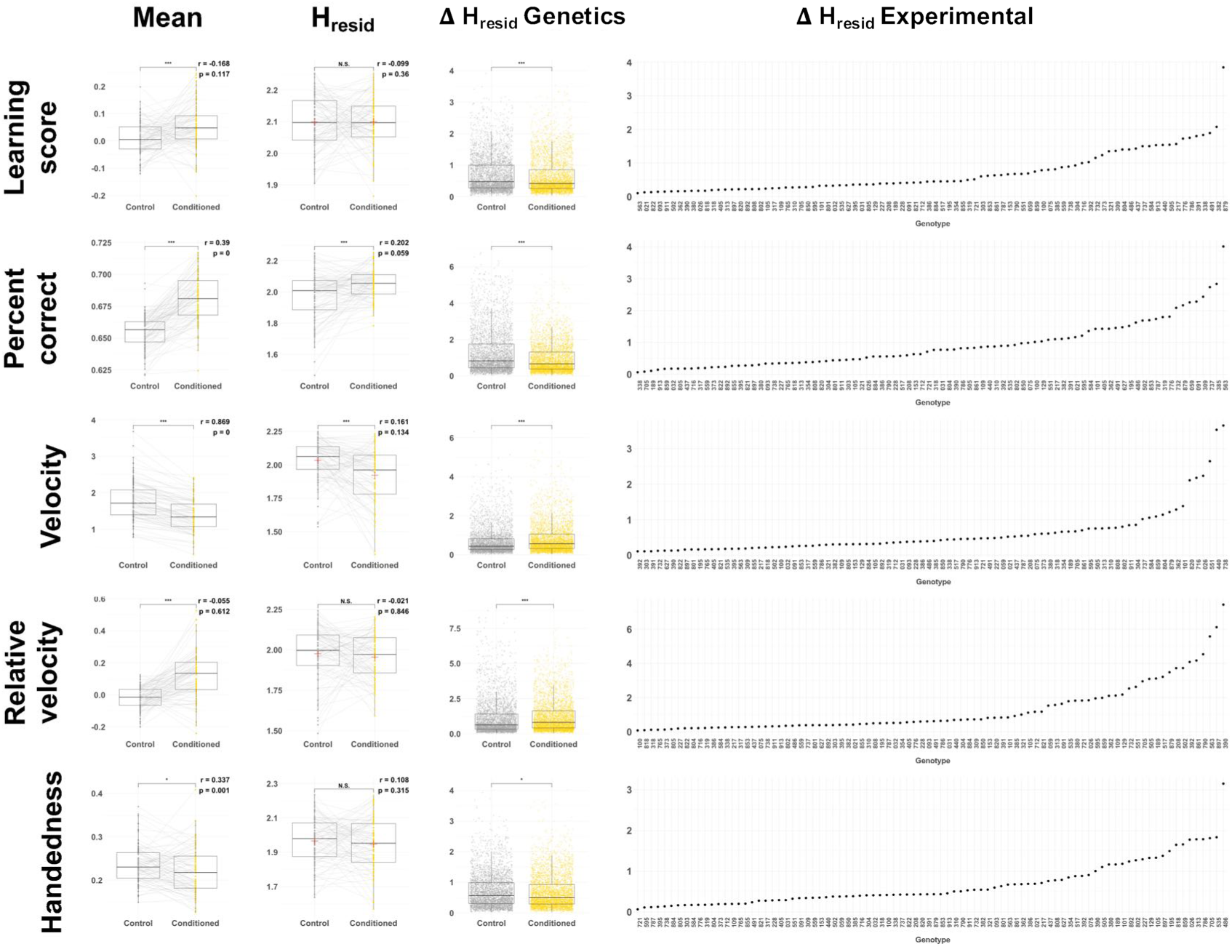
Phenotype means, residual entropies, Δ *H_resid_ Genetics* and Δ *H_resid_Experimental* (panels left to right) are shown for learning score, percent correctly made decisions, velocity, relative velocity and handedness (panels top to bottom). Δ *H_resid_ Experimental* is ranked smallest to largest, independently across phenotypes.

**Figure 2—figure supplement 8.**
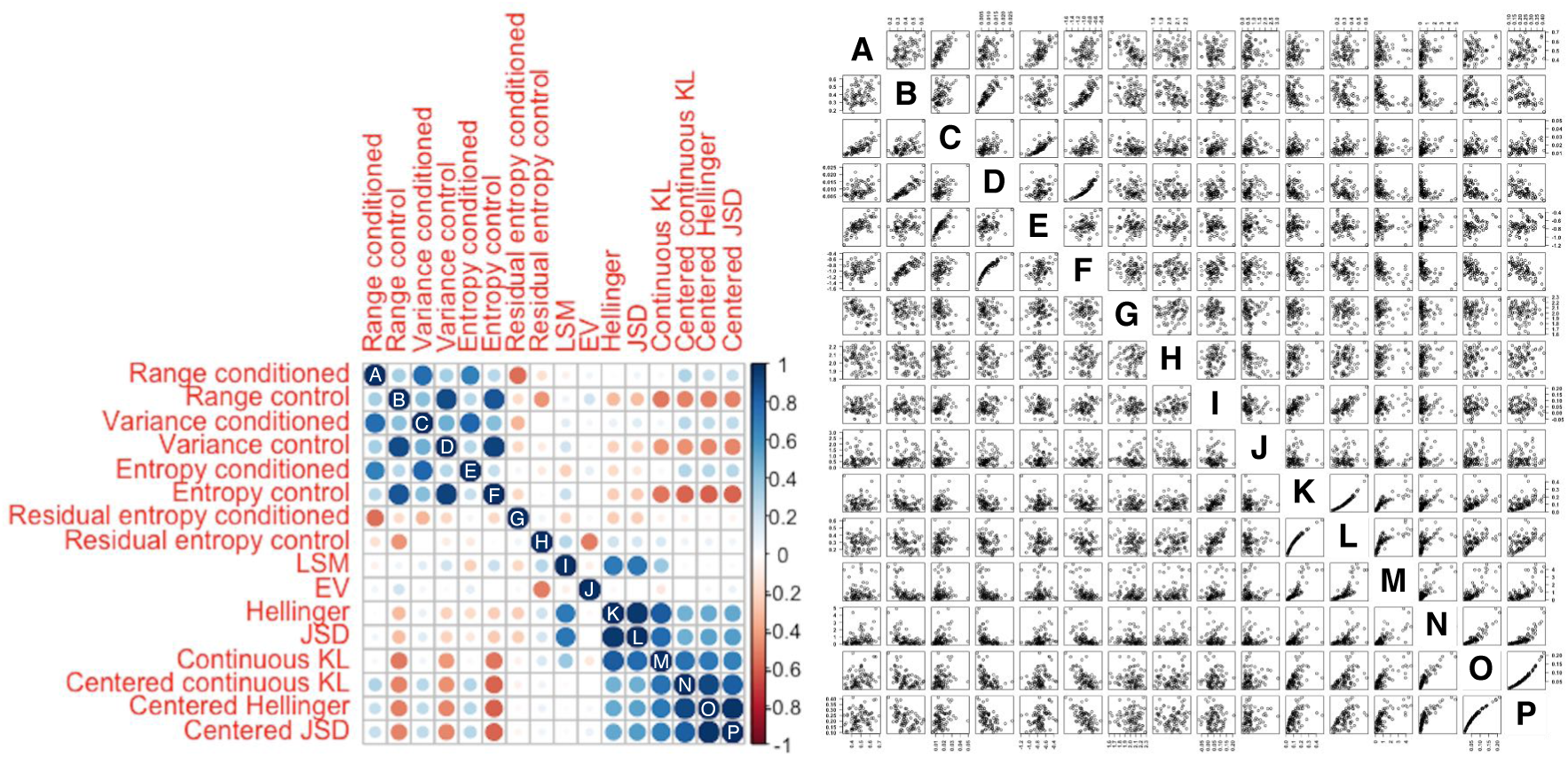
Measures of residual entropy are calculated for the two groups (control and conditioned) separately. Left panel is the correlation matrix heatmap, while the right panel shows the equivalent data scatterplots. Pearson product-moment correlation coefficient (r) scale: Blue r = 1, white r= 0, red r = -1.

**Figure 3—figure supplement 1.**
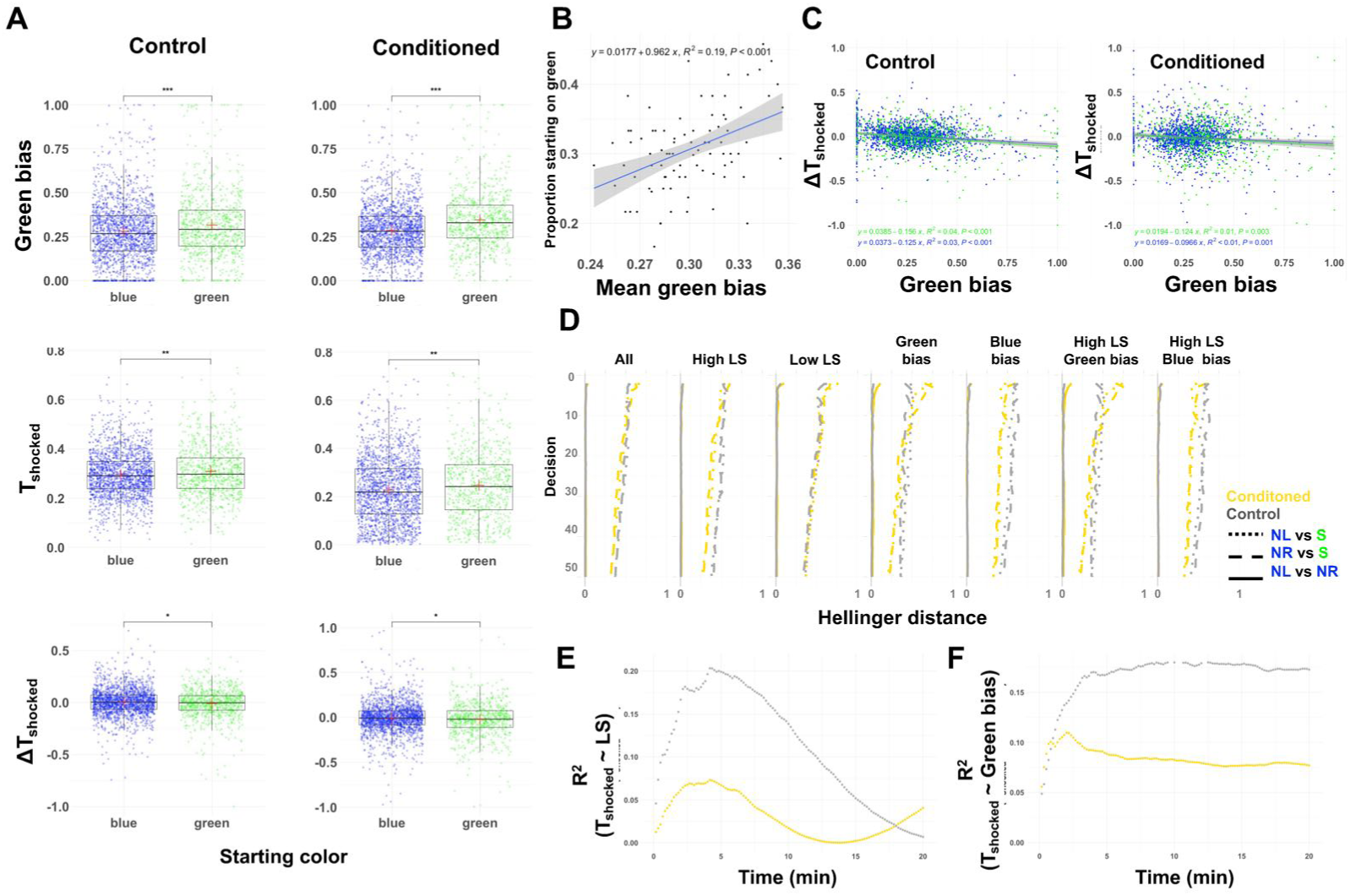
A) Starting position reflects colour bias and affects task performance. Flies with a higher green bias (top panels) tend to find themselves on the green arm at the beginning of the assay for both control (N = 2627, left) and conditioned (N = 2611, right) flies. This is later reflected in the task performance (middle panel) and change in task performance (bottom panel), where flies starting at green colour have higher T_shocked_ and lower change in T_shocked_ in green place learning. In other words, biased flies, and those that start the experiment on green colour are both disadvantaged and poorer learners. For all boxplots whiskers of boxplots show upper and lower quantiles, the bar indicates median across individuals (points). Asterisk indicates significance threshold for the T-test (* < 0.05,** < 0.01, *** < 0.001). N.S. indicates non-significance; B) Correlation between genotype’s green bias and proportion of flies on the green arm at the start of the conditioning for each genotype (N = 64 flies per genotype for 88 genotypes). Inset text show correlation equation, p-value and R^2^; C) Change in task performance with respect to green bias is shown for flies (points) that started the task on green (shown in green) or blue position in the Y-maze (shown in blue). Inset text show correlation equation, p-value and R^2^. While change in task performance is significantly dependent on the green bias, the slope of this association is not affected by starting position of the fly in either treatment. D) The evolution of Hellinger distance between distributions of cumulative task performance over 50 decisions of flies starting at different arms of the Y-maze (neutral left NL, neutral right NR and shocked S) is shown for conditioned (yellow) and control treatment (grey). Note that the distributions differ substantially and persistently depending on where the fly starts the assay. This was independent of the flies’ bias. The divergence between distributions is, however, reduced with learning. Distributions of T_shocked_ are calculated for flies that have made at least 50 decisions. E-F) Pearson correlation coefficient (R^2^) for correlation between cumulative task performance and learning score (E), and between cumulative task performance and green bias (F), over the time span of the conditioning. In all panels yellow indicates the conditioned and grey the control flies.

**Figure 3—figure supplement 2.**
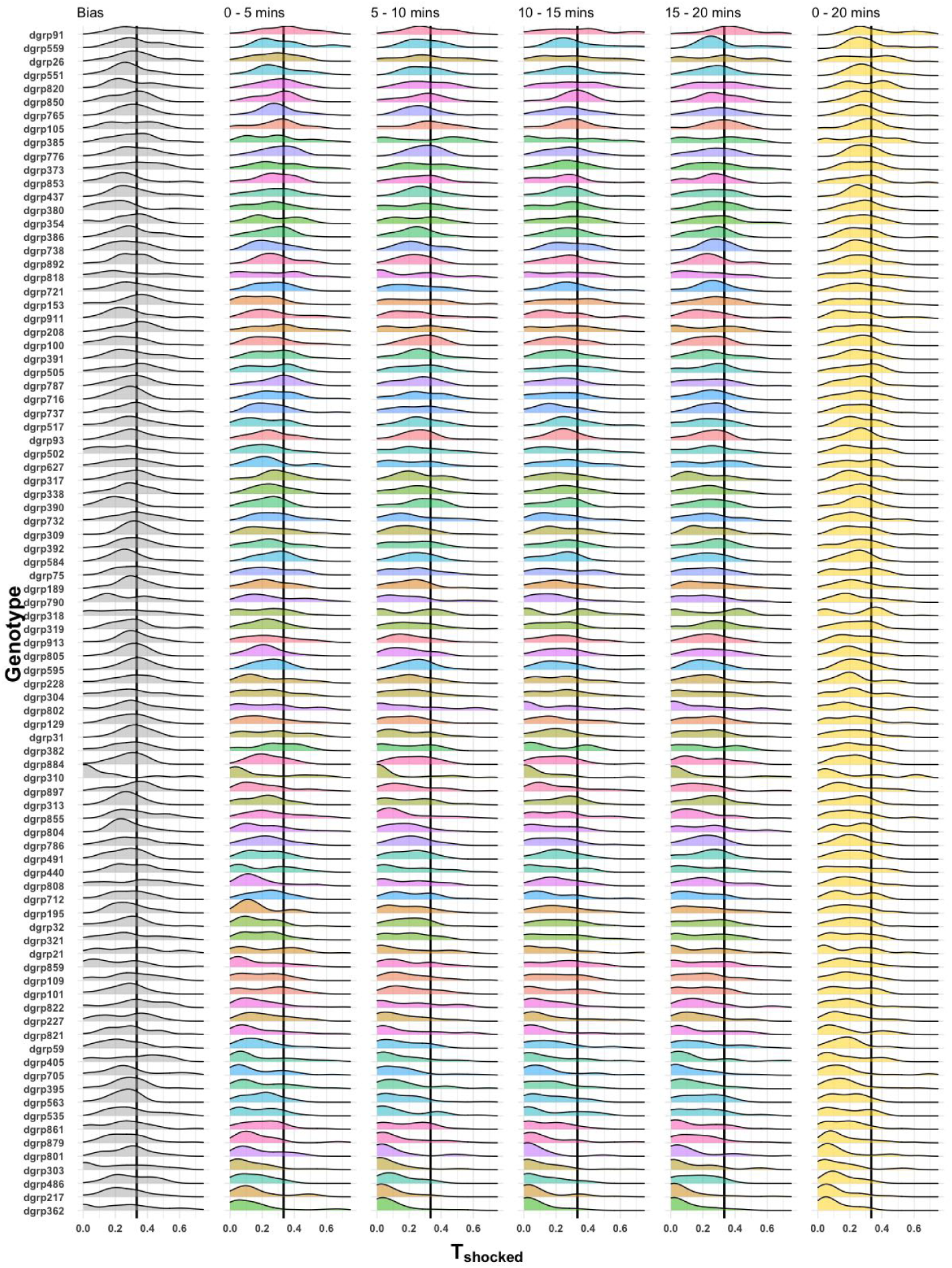
Bias estimated during the first 3 minutes of the assay (-3 mins to 0 mins), is shown in the first column. The 20 mins of the conditioning period after bias estimation is divided into 4 quarters of 5 minutes each, shown in the next 4 columns. The last panel shows the overall task performance during the 20 minutes of conditioning. The genotypes are ordered in rows in ascending order based on mean task performance of the conditioned flies.

**Figure 3—figure supplement 3.**
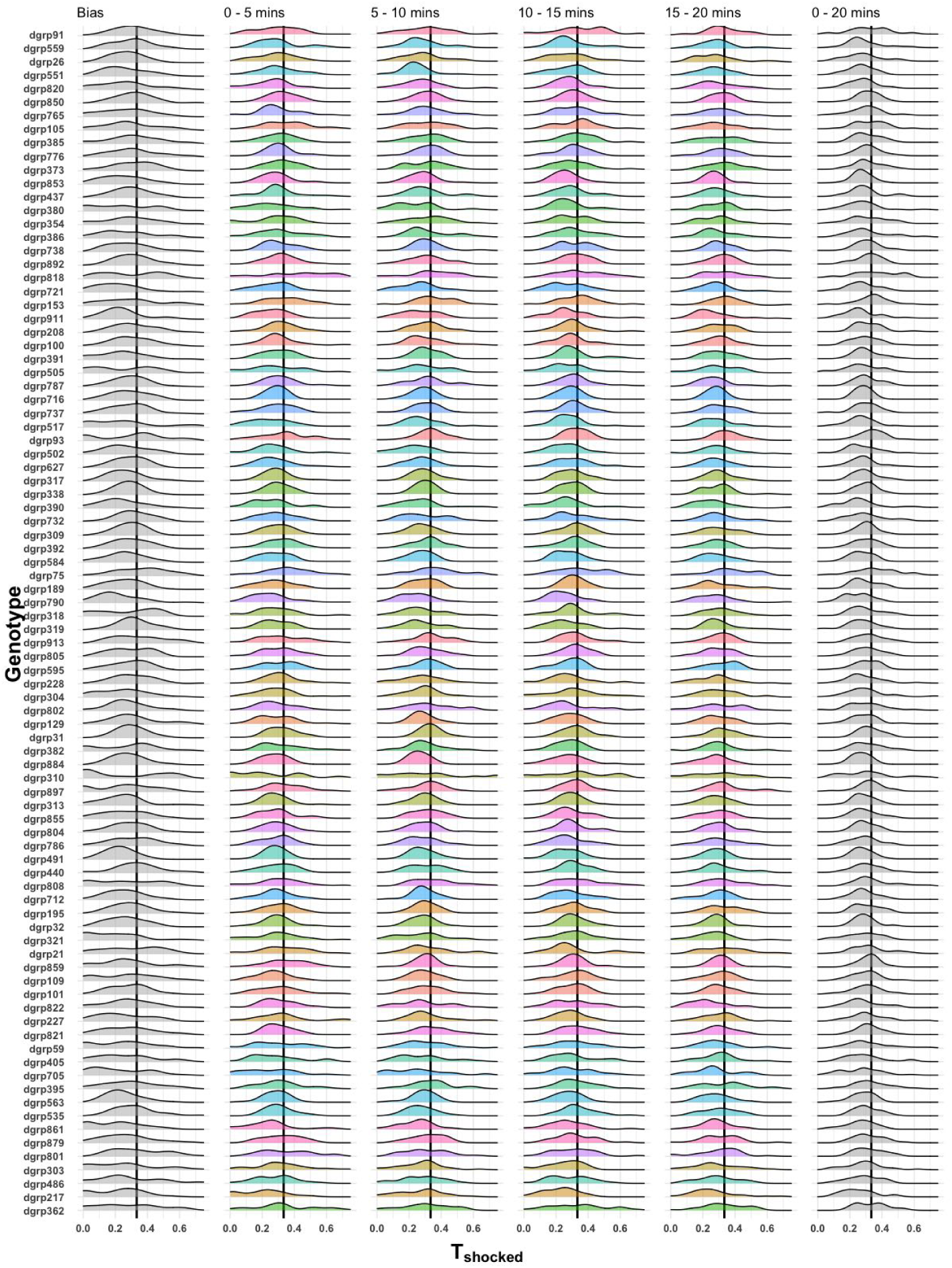
Bias estimated during the first 3 minutes of the assay (-3 mins to 0 mins), is shown in the first column. The 20 mins of the conditioning period after bias estimation is divided into 4 quarters of 5 minutes each, shown in the next 4 columns. The last panel shows the overall task performance during the 20 minutes of conditioning. The genotypes are ordered in ascending order of task performance of the conditioned flies (to match with Figure 3 - Figure supplement 3)

**Figure 4—figure supplement 1.**
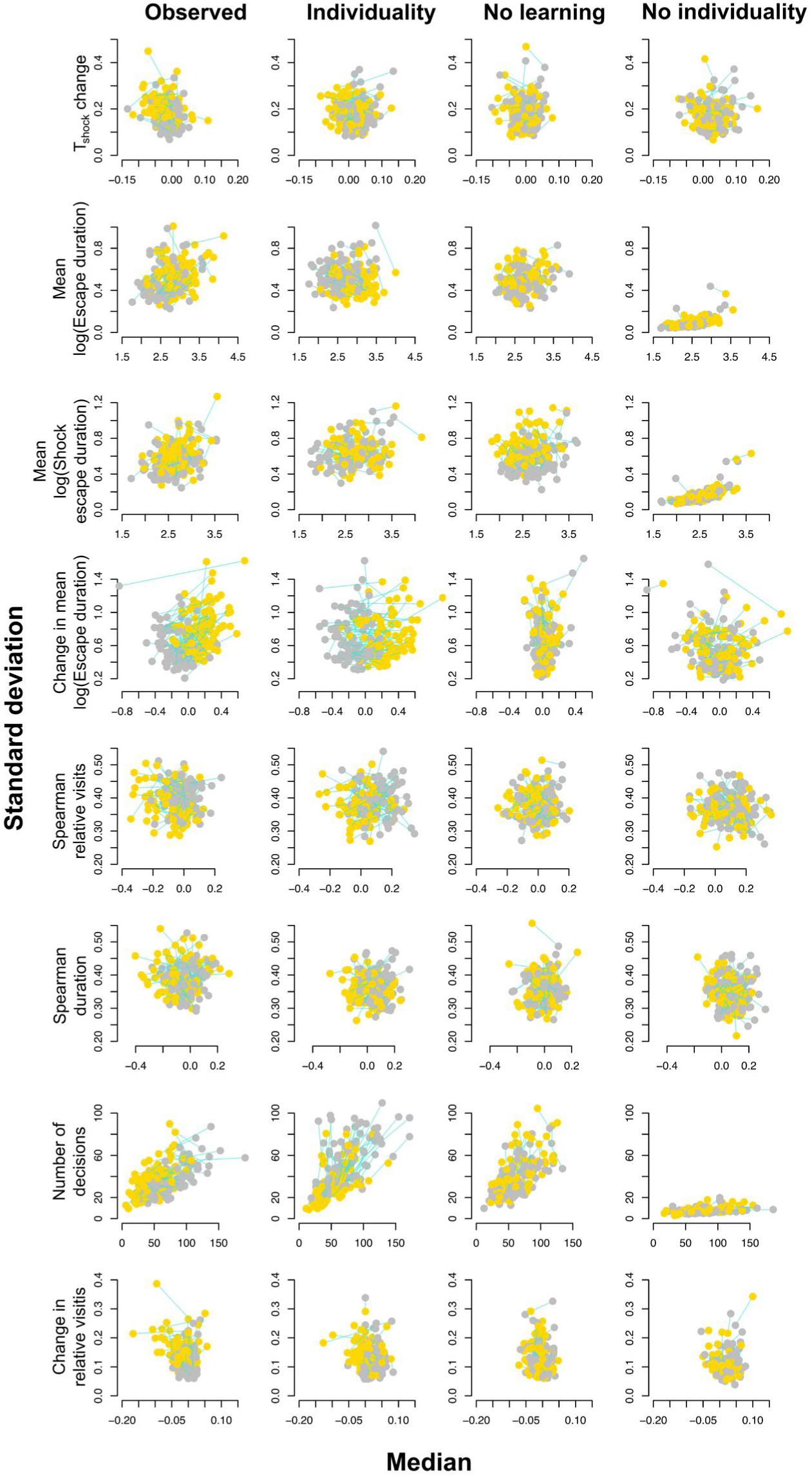
Points show genotype statistic in conditioned (yellow) and control group (grey). Behaviours shown are change in task performance, mean time to escape an arm, mean time to escape shock, change in time to escape an arm, Spearman’s rank correlation between four time intervals and relative number of visits to shocked arm (see Learning score in Methods), Spearman’s rank correlation between four time intervals and time spent in the shocked arm (see Learning score in Methods), number of made decisions and change the proportion of visits to the shocked arm. In all cases, the *Individuality* model provides the best representation of the observed data.

**Figure 4—figure supplement 2.**
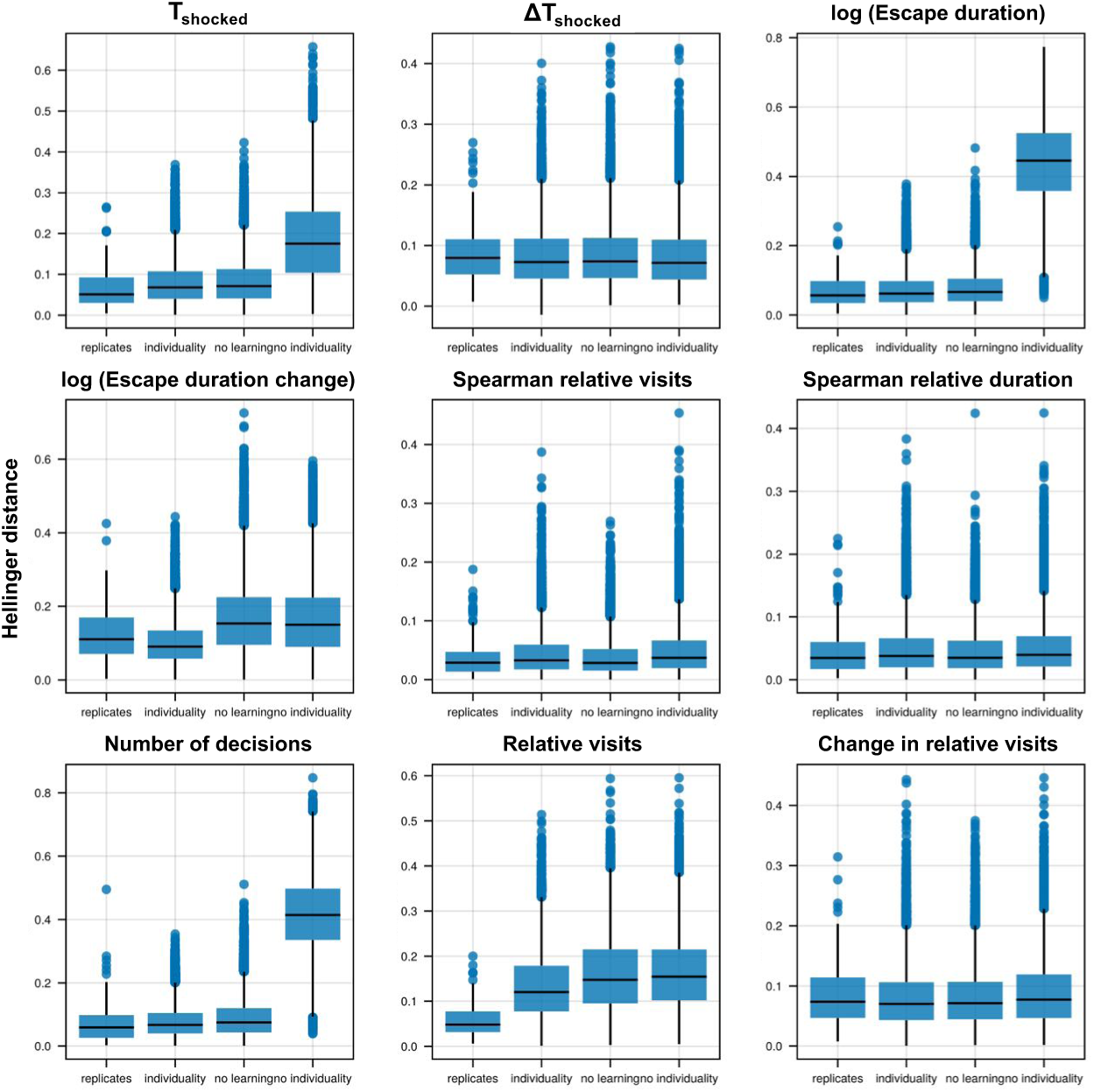
For each of the 9 behaviours, Hellinger distance is shown between observed data (N = 90 genotypes) and the equivalent data simulated 25 times based on three different models (*Individuality*, *No learning*, *No individuality*. Distance between observed and simulated replicates is also shown, where the first 32 simulated flies are compared to the first 32 observed flies (replicate 1) and the second 32 simulated to the second 32 observed (replicate 2). The *Individuality* model is able to reproduce the observed data, while *No individuality* model fails to capture the observed variation in the number of decisions, escape duration, change in escape duration, and the proportion of visits to shocked arm. Similarly, *No learning* model underperforms compared to *Individuality* model in capturing the change in escape duration and proportion of visits to shocked arm.

**Figure 4—figure supplement 3.**
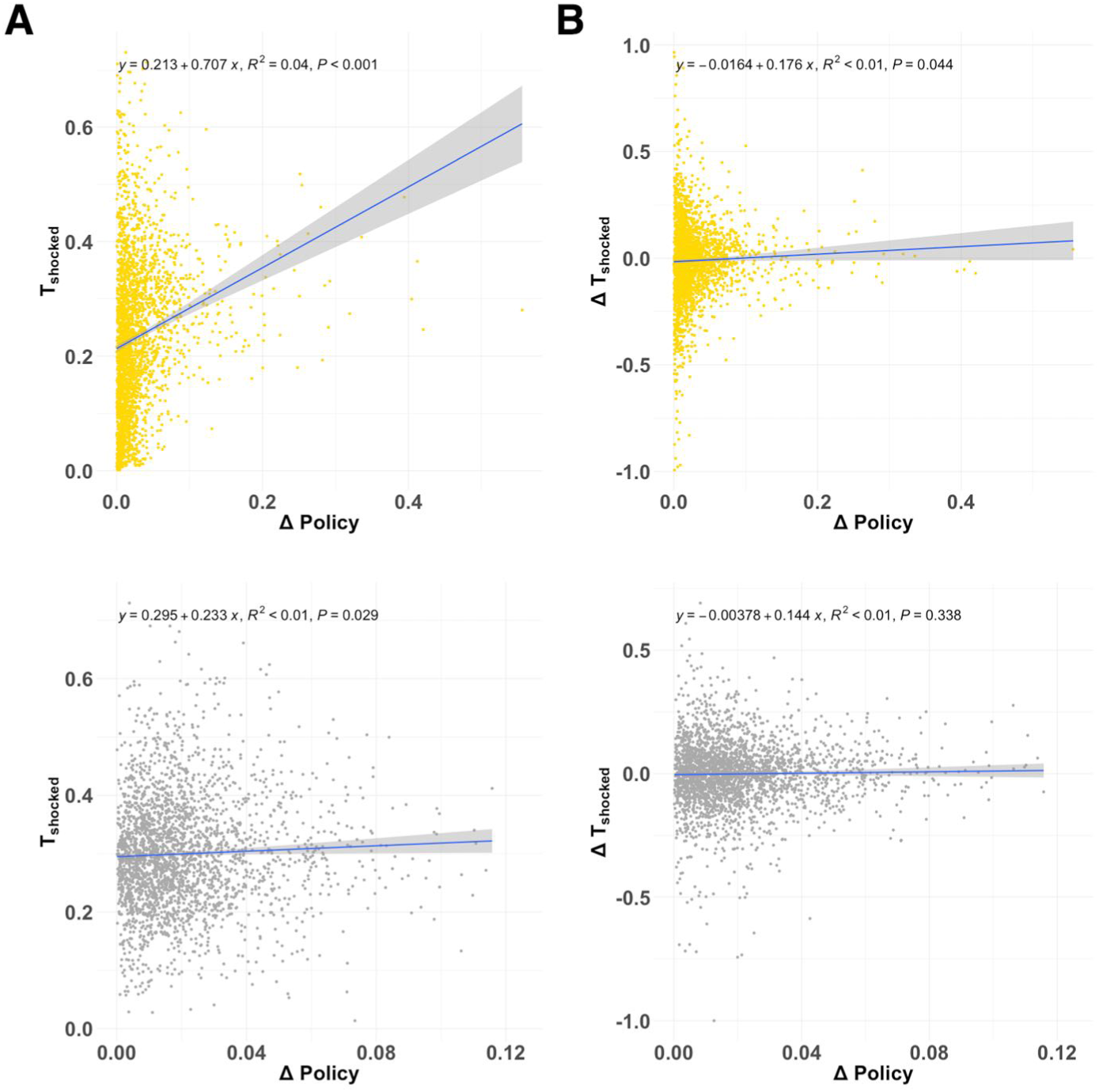
A) Task performance and B) change in task performance depend significantly on the change in behaviour policy in the conditioned treatment (yellow, upper panels). This dependence is much weaker or insignificant in the control treatment (grey, lower panels). Each point represents an individual fly. Inset text shows correlation equation, p-value and R^2^.

## References

Anderson C, Reiss I, Zhou C, Cho A, Siddiqi H, Mormann B, Avelis CM, Deford P, Bergland A, Roberts E, Taylor J, Vasiliauskas D, Johnston RJ. Natural variation in stochastic photoreceptor specification and color preference in Drosophila. eLife. 2017 Dec; 6:e29593. 10.7554/eLife.29593, doi: 10.7554/eLife.29593, publisher: eLife Sciences Publications, Ltd.

Arican C, Bulk J, Deisig N, Nawrot MP. Cockroaches Show Individuality in Learning and Memory During Classical and Operant Conditioning. Frontiers in Physiology. 2020; 10. https://www.frontiersin.org/journals/physiology/articles/10.3389/fphys.2019.01539.

Ayroles JF, Buchanan SM, O’Leary C, Skutt-Kakaria K, Grenier JK, Clark AG, Hartl DL, de Bivort BL. Behavioral idiosyncrasy reveals genetic control of phenotypic variability. Proceedings of the National Academy of Sciences. 2015 May; 112(21):6706–6711. 10.1073/pnas.1503830112, doi: 10.1073/pnas.1503830112, publisher: Proceedings of the National Academy of Sciences.

de Bivort B, Buchanan S, Skutt-Kakaria K, Gajda E, Ayroles J, O’Leary C, Reimers P, Akhund-Zade J, Senft R, Maloney R, Ho S, Werkhoven Z, Smith MAY. Precise Quantification of Behavioral Individuality From 80 Million Decisions Across 183,000 Flies. Frontiers in Behavioral Neuroscience. 2022; 16. https://www.frontiersin.org/articles/10.3389/fnbeh.2022.836626.

Boogert NJ, Madden JR, Morand-Ferron J, Thornton A. Measuring and understanding individual differences in cognition. Philosophical Transactions of the Royal Society B: Biological Sciences. 2018 Aug; 373(1756):20170280. 10.1098/rstb.2017.0280, doi: 10.1098/rstb.2017.0280, publisher: Royal Society.

Buchanan SM, Kain JS, de Bivort BL. Neuronal control of locomotor handedness in Drosophila. Proceedings of the National Academy of Sciences. 2015 May; 112(21):6700–6705. 10.1073/pnas.1500804112, doi: 10.1073/pnas.1500804112, publisher: Proceedings of the National Academy of Sciences.

Chen M, Sokolowski MB. How Social Experience and Environment Impacts Behavioural Plasticity in Drosophila. Fly. 2022 Dec; 16(1):68–84. 10.1080/19336934.2021.1989248, doi: 10.1080/19336934.2021.1989248, publisher: Taylor & Francis.

Chittka L, Dyer AG, Bock F, Dornhaus A. Bees trade off foraging speed for accuracy. Nature. 2003 Jul; 424(6947):388–388. 10.1038/424388a, doi: 10.1038/424388a.

Cover TM, Thomas JA. Elements of Information Theory. 2 ed. Hoboken, New Jersey.: John Wiley & Sons, Inc.; 2006.

Drost HG. Philentropy: Information Theory and Distance Quantification with R. Journal of Open Source Soft-ware. 2018 Jun; 3(26):765. http://joss.theoj.org/papers/10.21105/joss.00765, doi: 10.21105/joss.00765.

Dudai Y, Jan YN, Byers D, Quinn WG, Benzer S. dunce, a mutant of Drosophila deficient in learning. Proceedings of the National Academy of Sciences. 1976 May; 73(5):1684–1688. 10.1073/pnas.73.5.1684, doi: 10.1073/pnas.73.5.1684, publisher: Proceedings of the National Academy of Sciences.

Ernesto Salcedo, Armin Huber, Stefan Henrich, Linda V Chadwell, Wen-Hai Chou, Reinhard Paulsen, Steven G Britt. Blue- and Green-Absorbing Visual Pigments of *Drosophila*: Ectopic Expression and Physiological Characterization of the R8 Photoreceptor Cell-Specific Rh5 and Rh6 Rhodopsins. The Jour-nal of Neuroscience. 1999 Dec; 19(24):10716. http://www.jneurosci.org/content/19/24/10716.abstract, doi: 10.1523/JNEUROSCI.19-24-10716.1999.

Finke V, Baracchi D, Giurfa M, Scheiner R, Avarguès-Weber A. Evidence of cognitive specialization in an insect: proficiency is maintained across elemental and higher-order visual learning but not between sensory modalities in honey bees. Journal of Experimental Biology. 2021 Dec; 224(24):jeb242470. 10.1242/jeb.242470, doi: 10.1242/jeb.242470.

Frossard J, Renaud O. Permutation Tests for Regression, ANOVA, and Comparison of Signals: The permuco Package. Journal of Statistical Software. 2021 Oct; 99(15):1 – 32. https://www.jstatsoft.org/index.php/jss/article/view/v099i15, doi: 10.18637/jss.v099.i15, section: Articles.

Galsworthy MJ, Paya-Cano JL, Liu L, Monleón S, Gregoryan G, Fernandes C, Schalkwyk LC, Plomin R. Assessing Reliability, Heritability and General Cognitive Ability in a Battery of Cognitive Tasks for Laboratory Mice. Behavior Genetics. 2005 Sep; 35(5):675–692. 10.1007/s10519-005-3423-9, doi: 10.1007/s10519-005-3423-9.

Garrido-Jurado S, Muñoz-Salinas R, Madrid-Cuevas FJ, Marín-Jiménez MJ. Automatic generation and detection of highly reliable fiducial markers under occlusion. Pattern Recognition. 2014 Jun; 47(6):2280–2292. https://www.sciencedirect.com/science/article/pii/S0031320314000235, doi: 10.1016/j.patcog.2014.01.005.

Giurfa M, Zhang S, Jenett A, Menzel R, Srinivasan MV. The concepts of ‘sameness’ and ‘difference’ in an insect. Nature. 2001 Apr; 410(6831):930–933. 10.1038/35073582, doi: 10.1038/35073582.

Graham JH. Nature, Nurture, and Noise: Developmental Instability, Fluctuating Asymmetry, and the Causes of Phenotypic Variation. Symmetry. 2021; 13(7). doi: 10.3390/sym13071204.

Harden KP. Genetic determinism, essentialism and reductionism: semantic clarity for contested science. Nature Reviews Genetics. 2023 Mar; 24(3):197–204. 10.1038/s41576-022-00537-x, doi: 10.1038/s41576-022-00537-x.

Harris WA, Stark WS. Hereditary retinal degeneration in Drosophila melanogaster. A mutant defect associated with the phototransduction process. Journal of General Physiology. 1977 Mar; 69(3):261–291. 10.1085/jgp.69.3.261, doi: 10.1085/jgp.69.3.261.

Hausser J, Strimmer K. Entropy Inference and the James-Stein Estimator, With Application to Nonlinear Gene Association Networks. J Mach Learn Res. 2008 Nov; 10. doi: 10.1145/1577069.1755833.

Hellinger E. Neue Begründung der Theorie quadratischer Formen von unendlichvielen Veränderlichen. . 1909; 1909(136):210–271. 10.1515/crll.1909.136.210, doi: 10.1515/crll.1909.136.210.

Honegger K, de Bivort B. Stochasticity, individuality and behavior. Current Biology. 2018 Jan; 28(1):R8–R12. 10.1016/j.cub.2017.11.058, doi: 10.1016/j.cub.2017.11.058, publisher: Elsevier.

Hothorn T, Hornik K, van de Wiel MA, Zeileis A. Implementing a Class of Permutation Tests: The coin Package. Journal of Statistical Software. 2008 Nov; 28(8):1 – 23. https://www.jstatsoft.org/index.php/jss/article/view/v028i08, doi: 10.18637/jss.v028.i08, section: Articles.

J Lin. Divergence measures based on the Shannon entropy. IEEE Transactions on Information Theory. 1991 Jan; 37(1):145–151. doi: 10.1109/18.61115.

Kandel ER, Hawkins RD. The Biological Basis of Learning and Individuality. Scientific American. 1992; 267(3):78–87. http://www.jstor.org/stable/24939215, publisher: Scientific American, a division of Nature America, Inc.

Kempermann G, Lopes JB, Zocher S, Schilling S, Ehret F, Garthe A, Karasinsky A, Brandmaier AM, Lindenberger U, Winter Y, Overall RW. The individuality paradigm: Automated longitudinal activity tracking of large cohorts of genetically identical mice in an enriched environment. Neurobiology of Disease. 2022 Dec; 175:105916. https://www.sciencedirect.com/science/article/pii/S0969996122003084, doi: 10.1016/j.nbd.2022.105916.

Klingenberg CP. Phenotypic Plasticity, Developmental Instability, and Robustness: The Concepts and How They Are Connected. Frontiers in Ecology and Evolution. 2019; 7. https://www.frontiersin.org/journals/ecology-and-evolution/articles/10.3389/fevo.2019.00056.

Koellinger PD, Harden KP. Using nature to understand nurture. Science. 2018 Jan; 359(6374):386–387. 10.1126/science.aar6429, doi: 10.1126/science.aar6429, publisher: American Association for the Advancement of Science.

Korner AF. Individual differences at birth: Implications for early experience and later development. Amer-ican Journal of Orthopsychiatry. 1971; 41(4):608–619. doi: 10.1111/j.1939-0025.1971.tb03220.x, place: US Publisher: American Orthopsychiatric Association, Inc.

Kullback S, Leibler RA. On Information and Sufficiency. The Annals of Mathematical Statistics. 1951; 22(1):79–86. http://www.jstor.org/stable/2236703, publisher: Institute of Mathematical Statistics.

Laskowski KL, Bierbach D, Jolles JW, Doran C, Wolf M. The emergence and development of behavioral individuality in clonal fish. Nature Communications. 2022 Oct; 13(1):6419. 10.1038/s41467-022-34113-y, doi: 10.1038/s41467-022-34113-y.

Lin WC, Delevich K, Wilbrecht L. A role for adaptive developmental plasticity in learning and decision making. Sensitive and critical periods. 2020 Dec; 36:48–54. https://www.sciencedirect.com/science/article/pii/S2352154620301121, doi: 10.1016/j.cobeha.2020.07.010.

Linneweber GA, Andriatsilavo M, Dutta SB, Bengochea M, Hellbruegge L, Liu G, Ejsmont RK, Straw AD, Wernet M, Hiesinger PR, Hassan BA. A neurodevelopmental origin of behavioral individuality in the Drosophila visual system. Science. 2020 Mar; 367(6482):1112–1119. 10.1126/science.aaw7182, doi: 10.1126/science.aaw7182, publisher: American Association for the Advancement of Science.

Little CC. Variability and Individuality. Science. 1933; 77(1990):195–197. http://www.jstor.org/stable/1658280, publisher: American Association for the Advancement of Science.

Luo L, Sun T, Guan X, Ni Y, Yang L, Zhao Q, Kong X, Chen Y, Zhang J. Advanced Parental Age Impaired Fear Conditioning and Hippocampal LTD in Adult Female Rat Offspring. Neurochemical Research. 2017 Oct; 42(10):2869–2880. 10.1007/s11064-017-2306-9, doi: 10.1007/s11064-017-2306-9.

Mackay TFC, Richards S, Stone EA, Barbadilla A, Ayroles JF, Zhu D, Casillas S, Han Y, Magwire MM, Cridland JM, Richardson MF, Anholt RRH, Barrón M, Bess C, Blankenburg KP, Carbone MA, Castellano D, Chaboub L, Duncan L, Harris Z, et al. The Drosophila melanogaster Genetic Reference Panel. Nature. 2012 Feb; 482(7384):173–178. 10.1038/nature10811, doi: 10.1038/nature10811.

Maloney R, Ye A, Saint-Pre SK, Alisch T, Zimmerman D, Pittoors N, de Bivort BL. Drift in Individual Behavioral Phenotype as a Strategy for Unpredictable Worlds. bioRxiv. 2024 Jan; p. 2024.09.05.611301. http://biorxiv.org/content/early/2024/09/10/2024.09.05.611301.abstract, doi: 10.1101/2024.09.05.611301.

Mao WJ, Wu ZY, Yang ZH, Xu YW, Wang SQ. Advanced maternal age impairs spatial learning capacity in young adult mouse offspring. . 2018; 10(3):975–988.

Martin GM. Epigenetic drift in aging identical twins. Proceedings of the National Academy of Sciences. 2005 Jul; 102(30):10413–10414. 10.1073/pnas.0504743102, doi: 10.1073/pnas.0504743102, publisher: Proceedings of the National Academy of Sciences.

Melnattur KV, Pursley R, Lin TY, Ting CY, Smith PD, Pohida T, Lee CH. Multiple Redundant Medulla Projection Neurons Mediate Color Vision in Drosophila. Journal of Neurogenetics. 2014 Dec; 28(3-4):374–388. 10.3109/01677063.2014.891590, doi: 10.3109/01677063.2014.891590, publisher: Taylor & Francis.

Mery F, Burns JG. Behavioural plasticity: an interaction between evolution and experience. Evolutionary Ecol-ogy. 2010 May; 24(3):571–583. 10.1007/s10682-009-9336-y, doi: 10.1007/s10682-009-9336-y.

Modi MN, Rajagopalan AE, Rouault H, Aso Y, Turner GC. Flexible specificity of memory in Drosophila depends on a comparison between choices. eLife. 2023 Jun; 12:e80923. 10.7554/eLife.80923, doi: 10.7554/eLife.80923, publisher: eLife Sciences Publications, Ltd.

Mollá-Albaladejo R, Sánchez-Alcañiz JA. Behavior Individuality: A Focus on Drosophila melanogaster. Frontiers in Physiology. 2021; 12. https://www.frontiersin.org/articles/10.3389/fphys.2021.719038.

Nepoux V, Babin A, Haag C, Kawecki TJ, Le Rouzic A. Quantitative genetics of learning ability and resistance to stress in Drosophila melanogaster. Ecology and Evolution. 2015 Feb; 5(3):543–556. 10.1002/ece3.1379, doi: 10.1002/ece3.1379, publisher: John Wiley & Sons, Ltd.

Nouvian M, Galizia CG. Aversive Training of Honey Bees in an Automated Y-Maze. Frontiers in Physiology. 2019; 10. https://www.frontiersin.org/articles/10.3389/fphys.2019.00678.

Partridge L, Sgrò CM. Behavioural genetics: Molecular genetics meets feeding ecology. Current Biology. 1998 Jan; 8(1):R23–R24. 10.1016/S0960-9822(98)70011-9, doi: 10.1016/S0960-9822(98)70011-9, publisher: Elsevier.

Álvarez Quintero N, Kim SY. Effects of maternal age and environmental enrichment on learning ability and brain size. Behavioral Ecology. 2024 Jul; 35(4):arae049. 10.1093/beheco/arae049, doi: 10.1093/beheco/arae049.

Rozenfeld E, Parnas M. Neuronal circuit mechanisms of competitive interaction between action-based and coincidence learning. Science Advances. 2024; 10(49):eadq3016. 10.1126/sciadv.adq3016, doi: 10.1126/sciadv.adq3016.

Schnaitmann C, Garbers C, Wachtler T, Tanimoto H. Color Discrimination with Broadband Photoreceptors. Current Biology. 2013 Dec; 23(23):2375–2382. https://www.sciencedirect.com/science/article/pii/S0960982213013122, doi: 10.1016/j.cub.2013.10.037.

Shannon CE. A Mathematical Theory of Communication. Bell System Technical Journal. 1948 Jul; 27(3):379–423. 10.1002/j.1538-7305.1948.tb01338.x, doi: 10.1002/j.1538-7305.1948.tb01338.x, publisher: John Wiley & Sons, Ltd.

Smith MAY, Honegger KS, Turner G, de Bivort B. Idiosyncratic learning performance in flies. Biology Letters. 2022 Feb; 18(2):20210424. 10.1098/rsbl.2021.0424, doi: 10.1098/rsbl.2021.0424, publisher: Royal Society.

Smith P, Arias R, Sonti S, Odgerel Z, Santa-Maria I, McCabe BD, Tsaneva-Atanasova K, Louis ED, Hodge JJL, Clark LN. A Drosophila Model of Essential Tremor. Scientific Reports. 2018 May; 8(1):7664. 10.1038/s41598-018-25949-w, doi: 10.1038/s41598-018-25949-w.

Sridhar VH, Li L, Gorbonos D, Nagy M, Schell BR, Sorochkin T, Gov NS, Couzin ID. The geometry of decision-making in individuals and collectives. Proceedings of the National Academy of Sciences. 2021 Dec; 118(50):e2102157118. http://www.pnas.org/content/118/50/e2102157118.abstract, doi: 10.1073/pnas.2102157118.

Strasser H, Weber C. On the asymptotic theory of permutation statistics. Report, Vienna University of Eco-nomics and Business Administration; 1999. https://books.google.ch/books?id=pieBNAEACAAJ.

Thomas F Mathejczyk, Cara Knief, Muhammad A Haidar, Florian Freitag, Tydings McClary, Mathias F Wernet, Gerit A Linneweber. Individuality across environmental context in D *rosophila melanogaster*. bioRxiv. 2023 Jan; p. 2023.11.26.568741. http://biorxiv.org/content/early/2023/11/26/2023.11.26.568741. abstract, doi: 10.1101/2023.11.26.568741.

Tosh CR, Brogan B. Environmental diversity constrains learning in Drosophila melanogaster. Ecological En-tomology. 2017 Dec; 42(6):697–703. 10.1111/een.12435, doi: 10.1111/een.12435, publisher: John Wiley & Sons, Ltd.

Turner CH. The homing of ants: an experimental study of ant behavior. University of Chicago.; 1907.

Turner CH. Experiments on pattern-vision of the honey bee. The Biological Bulletin. 1911; 21(5):249–264. Publisher: Marine Biological Laboratory.

Wang X, Amei A, de Belle JS, Roberts SP. Environmental effects on Drosophila brain development and learn-ing. Journal of Experimental Biology. 2018 Jan; 221(1):jeb169375. 10.1242/jeb.169375, doi: 10.1242/jeb.169375.

Whishaw IQ. Place Learning in Hippocampal Rats and the Path Integration Hypothesis. Neuroscience & Biobehavioral Reviews. 1998 Mar; 22(2):209–220. https://www.sciencedirect.com/science/article/pii/S014976349700002X, doi: 10.1016/S0149-7634(97)00002-X.

Williams-Simon PA, Posey C, Mitchell S, Ng’oma E, Mrkvicka JA, Zars T, King EG. Multiple genetic loci affect place learning and memory performance in Drosophila melanogaster. Genes, Brain and Behavior. 2019 Sep; 18(7):e12581. 10.1111/gbb.12581, doi: 10.1111/gbb.12581, publisher: John Wiley & Sons, Ltd.

Xu PS, Lee D, Holy TE. Experience-Dependent Plasticity Drives Individual Differences in Pheromone-Sensing Neurons. Neuron. 2016 Aug; 91(4):878–892. https://www.sciencedirect.com/science/article/pii/S0896627316304196, doi: 10.1016/j.neuron.2016.07.034.

